# Dynamic top-down biasing implements rapid adaptive changes to individual movements

**DOI:** 10.1101/2022.06.16.496455

**Authors:** Lucas Y Tian, Timothy L. Warren, Michael S. Brainard

## Abstract

Complex behaviors depend on the coordinated activity of neural ensembles in interconnected brain areas. The behavioral function of such coordination, often measured as co-fluctuations in neural activity across areas, is poorly understood. One hypothesis is that rapidly varying co-fluctuations may be a signature of moment-by-moment task-relevant influences of one area on another. We tested this possibility for error-corrective adaptation of birdsong, a form of motor learning which has been hypothesized to depend on the top-down influence of a higher-order area, LMAN, in shaping moment-by-moment output from a primary motor area, RA. In paired recordings of LMAN and RA in singing birds, we discovered a neural signature of a top-down influence of LMAN on RA, quantified as an LMAN-leading co-fluctuation between these areas. During learning, this cofluctuation strengthened in a premotor temporal window linked to the specific movement, sequential context, and acoustic modification associated with learning. Moreover, transient perturbation of LMAN activity specifically within this premotor window caused rapid occlusion of pitch modifications, consistent with LMAN conveying a temporally localized motor-biasing signal. Combined, our results reveal a dynamic top-down influence of LMAN on RA that varies on the rapid timescale of individual movements and is flexibly linked to contexts associated with learning. This finding indicates that inter-area co-fluctuations can be a signature of dynamic top-down influences that support complex behavior and its adaptation.

## Introduction

Complex behaviors depend on the coordinated activity of neural ensembles in distinct brain areas, yet much remains to be resolved about the specific behavioral functions of such multi-region coordination. Recent analyses of trial-by-trial activity co-fluctuations among neuronal populations in connected brain regions have noted a link between changes to coordinated activity and behavioral performance (Cowley et al., 2020; Koralek et al., 2013, 2012; Lemke et al., 2019; Makino et al., 2017; Perich et al., 2018; Sawada et al., 2015; Veuthey et al., 2020; Wagner et al., 2019). Although some slower activity co-fluctuations may reflect task-independent, shared changes in a global internal state (Cowley et al., 2020), more dynamic co-fluctuations—varying rapidly on a moment-by-moment basis and linked flexibly to specific behavioral contexts—could be a signature of task-relevant influences of one area on another (Semedo et al., 2019).

One intriguing role for such dynamic inter-area influences has been hypothesized in the context of the learning and adaptation of skilled reaching behavior (Perich et al., 2018; Veuthey et al., 2020). Specifically, the joint evolution of inter-area activity co-fluctuations and behavioral performance over the course of learning has led to the suggestion that top-down inputs from higher-order areas dynamically implement learning-related changes in activity in primary motor areas. In principle, such top-down modulation during learning could operate to control adaptive behavioral modifications on the fine time scale of individual movements. However, the difficulty in identifying and quantifying discrete movement components during skilled reaching leaves unresolved whether such top-down influences operate on this rapid time scale during learning.

Moment-by-moment shaping of activity in primary motor areas by dynamic, top-down control has been hypothesized to instantiate error-corrective adaptation of birdsong—a complex learned behavior with quantifiable, precisely controlled and rapidly varying movements. Indeed, previous work has demonstrated that pharmacologically inactivating either the song-specific higher-order area LMAN or its synapses in the primary motor cortical analog RA (Fig. 1A) transiently eliminates recently learned modifications to pitch (fundamental frequency) of the individual 50-200 ms syllables that constitute the adult song [Fig. 1B, (Andalman and Fee, 2009; Charlesworth et al., 2012; Tian and Brainard, 2017; Warren et al., 2011)]. This raises the possibility that LMAN provides top-down drive to the motor pathway to implement adaptive changes to behavior (Ali et al., 2013; Andalman and Fee, 2009; Aronov et al., 2008; Bottjer et al., 1984; Charlesworth et al., 2012; Doya and Sejnowski, 1998; Fee and Goldberg, 2011; Kao et al., 2005; Kearney et al., 2019; Nordeen and Nordeen, 2010; Olveczky et al., 2005; Scharff and Nottebohm, 1991; Tian and Brainard, 2017; Troyer and Bottjer, 2001; Troyer and Doupe, 2000a; Warren et al., 2011). Indeed, studies in sleeping and anesthetized birds have identified co-fluctuations in LMAN and RA activity (Hahnloser et al., 2006; Kimpo et al., 2003)—measured as increases in the cross-covariance between LMAN and RA activity—that peak at a short LMAN-leading temporal lag, consistent with the possibility of a dynamic top-down influence of LMAN on RA. However, it remains unclear whether such LMAN-RA co-fluctuations are present during singing and, if so, whether they change during learning in a manner that could support a top-down role of LMAN in the adaptive adjustment of song parameters. These observations, together with the specificity and temporal precision with which adult song can be modified, render birdsong an ideal system for testing whether changes to inter-area activity co-fluctuations develop with the appropriate specificity to constitute top-down commands for implementing behavioral modifications on the time scale of individual movements.

**Figure 1.**
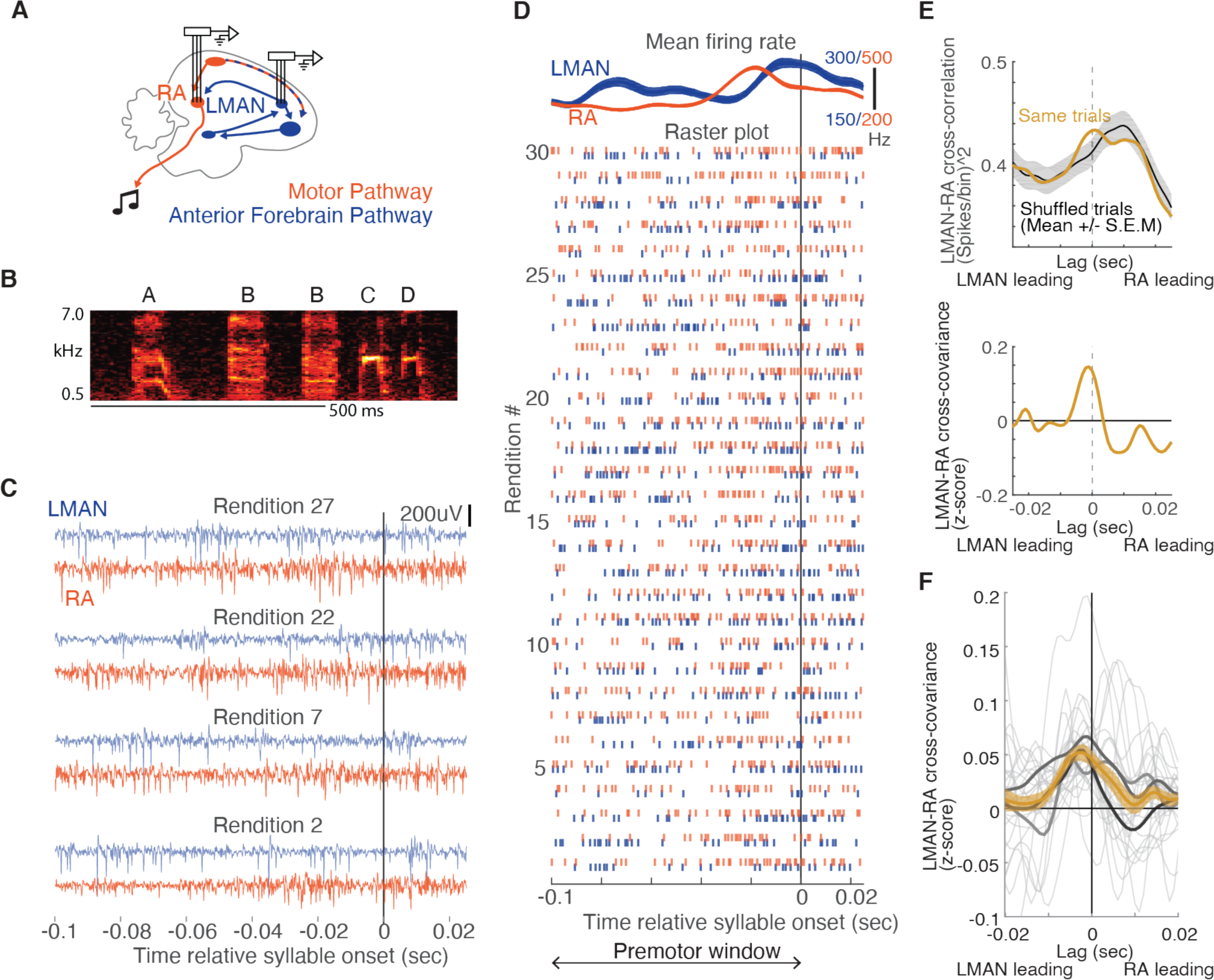
LMAN-leading co-fluctuations of LMAN and RA activity are present during singing. (A) Schematic of song system circuitry implicated in the production and learning of birdsong. This study focuses on hypothesized top-down signals from LMAN (lateral magnocellular nucleus of the anterior nidopallium), the output nucleus of the Anterior Forebrain Pathway (blue), to RA (robust nucleus of the arcopallium) in the Motor Pathway (red). Recordings were made using multi-elec-trode arrays chronically implanted in LMAN and RA. (B) Spectrogram of a single bout of a sequence of syllables forming a motif labeled “ABBCD”. A motif is a specific sequence of syllables that is consistently sung across bouts and is unique to a bird (see Methods). Scale bar, 500 ms. (C) Example paired recordings in LMAN and RA across four renditions (numbered to match those in panel D) aligned to the onset of a single syllable, filtered in the spike band (300-3000 Hz). (D) Raster plot representing spikes for the same pair of LMAN and RA sites shown in panel C, across thirty renditions, ordered chronologically and aligned to syllable onset. Above the raster plot are the mean (+/- SEM) smoothed firing rates. “Premotor window” refers to the time period of neural activity used for calculating the cross-covariance between LMAN and RA spike trains for each syllable. (E) Calculation of normalized cross-covariance for the example sites and syllable in panels C and D. Top: cross-correlation was calculated using spike trains for both concurrent trials (“Same trials”) and a control dataset in which LMAN and RA activity were shuffled between adjacent renditions (“Shuffled trials”). We used this shuffling to estimate the cross-correlation between the mean activity patterns in LMAN and RA, see Methods). Bottom: the mean of the shuffle-computed crosscorrelation functions (“Shuffled trials”) was subtracted from the actual cross-correlation function (“Same trials”) to compute cross-covariance. We then divided the cross-covariance by the standard deviation of the shuffled cross-correlations to compute a normalized cross-covariance (measured in z-score). (F) LMAN-RA cross-covariance across all syllables and pairs of LMAN and RA sites during baseline singing. The light grey curves represent individual syllables, averaged over all simultaneously recorded pairs of LMAN and RA sites. The dark grey curves represent individual birds (N = 3), averaged across syllables. The orange curve represents the mean +/S.E.M. across all syllables (N=27).

## Results

### LMAN-leading co-fluctuations of LMAN and RA activity are present during singing

To determine whether the co-fluctuations in neural activity between LMAN and RA previously identified in sleeping and anaesthetized birds are present within the premotor window for individual syllables, we first examined neural activity in both nuclei during baseline singing before the onset of training (Fig. S1; multi-unit recordings, N=30 LMAN and 52 RA sites, 4 birds). Consistent with prior studies in which LMAN and RA were recorded in separate singing birds (Aronov et al., 2008; Chi and Margoliash, 2001; Hessler and Doupe, 1999; Kao et al., 2008; Leonardo and Fee, 2005; Margoliash and Yu, 1996; McCasland, 1987; Olveczky et al., 2005; Sober et al., 2008), average activity in both areas exhibited consistent temporal structure aligned to ongoing song, peaking within 50 ms prior to syllable onsets (Fig. S1C,D). Additionally, we observed a close alignment between the patterns of activity in the two areas (Fig. S1C, E, F). Despite such consistent temporal structure in average activity, we observed rendition-by-rendition variation in the patterning of activity in both nuclei, raising the possibility that this variation reflects cofluctuations in LMAN and RA activity. We assessed LMAN-RA co-fluctuation within a premotor window for each syllable, which we defined as the 100 ms period directly preceding syllable onset, chosen based on empirical estimates of when LMAN and RA activity patterns maximally influence acoustic structure of the upcoming syllable (Fee et al., 2004; Giret et al., 2014; Kao et al., 2005; Kojima et al., 2018; Margoliash and Yu, 1996; Sober et al., 2008).

We assessed co-fluctuations of LMAN and RA activity by measuring cross-covariance, a metric previously used to uncover LMAN-leading co-fluctuations in sleeping and anaesthetized birds [(Kimpo et al., 2003), see Methods]. For a given pair of LMAN and RA recordings (example in Fig. 1C-F), LMAN-RA cross-covariance is a measure of the similarity of moment-by-moment fluctuations of LMAN and RA spike patterns away from their respective means, computed as a function of the time lag between these patterns (Perkel et al., 1967). In birds singing spontaneously at baseline (“undirected” singing produced in isolation), the LMAN-RA cross-covariance was on average positive during the premotor window for individual syllables (example pair of sites in Fig. 1E; summary across pairs of recordings and syllables in Fig. 1F), similar to cross-covariance measurements in anesthetized and sleeping birds (Hahnloser et al., 2006; Kimpo et al., 2003). Although there was variation in the magnitude and time lag of cross-covariance peaks across syllables and recording sites (see light gray curves in Fig. 1F) there was a dominant positive peak in the average cross-covariance with LMAN leading at a short time lag [∼3 ms, Fig. 1F; c.f. (Kimpo et al., 2003)], This putative influence of LMAN on RA during baseline singing is consistent with the known presence of a direct excitatory projection from LMAN to RA (Bottjer et al., 1989; Kubota and Saito, 1991; Mooney and Konishi, 1991; Nottebohm et al., 1982), and a hypothesized role of LMAN in driving variability in the motor pathway for trial-and-error learning (Giret et al., 2014; Kao et al., 2005; Kao and Brainard, 2006; Kojima et al., 2018; Olveczky et al., 2005; Stepanek and Doupe, 2010), and raises the question of whether similar top-down signals from LMAN to RA operate in the context of learning to instantiate rapid adaptive adjustments to individual syllables.

### LMAN-RA co-fluctuations are enhanced during learning

To determine whether adaptive biasing of individual syllables during learning is accompanied by changes to co-fluctuations in LMAN and RA activity, we examined LMAN-RA crosscovariance while birds learned modifications to the pitch of individual song syllables in response to pitch-contingent aversive white noise (WN) feedback (Fig. 2A, B) (Ali et al., 2013; Andalman and Fee, 2009; Charlesworth et al., 2012, 2011; Tian and Brainard, 2017; Tumer and Brainard, 2007; Warren et al., 2011). On average, across syllables targeted for pitch modification, the LMAN-RA cross-covariance was enhanced following training (3 to 10 hours) with performance-contingent reinforcement (Fig. 2A-D, F, “Target syllable”; cross-covariance function in Fig. 2D; summary histogram using the area under the curve in Fig. 2F) and was maximal at a short LMAN-leading time lag (∼2 ms, Fig. 2D). Moreover, LMAN-RA cross-covariance increased with a similar time course to pitch modification (Fig. 2G, H). In contrast, there were no detectable changes in the cross- covariance for non-targeted syllables (Fig. 2E, F, “Non-target syllables”), including those directly preceding the targeted syllable (Fig. S2A). There was also no detectable learning-related difference between targeted and non-targeted syllables in changes in the overall level of activity within each nucleus (Fig. S2B). The observations that increases in LMAN-RA cross-covariance are restricted to the targeted syllable and develop in parallel with pitch modification argue that these changes are linked specifically to learning, and not to other processes such as changes to arousal, non- stationarity or “drift” of recorded units, or circadian variation in neural activity patterns.

**Figure 2.**
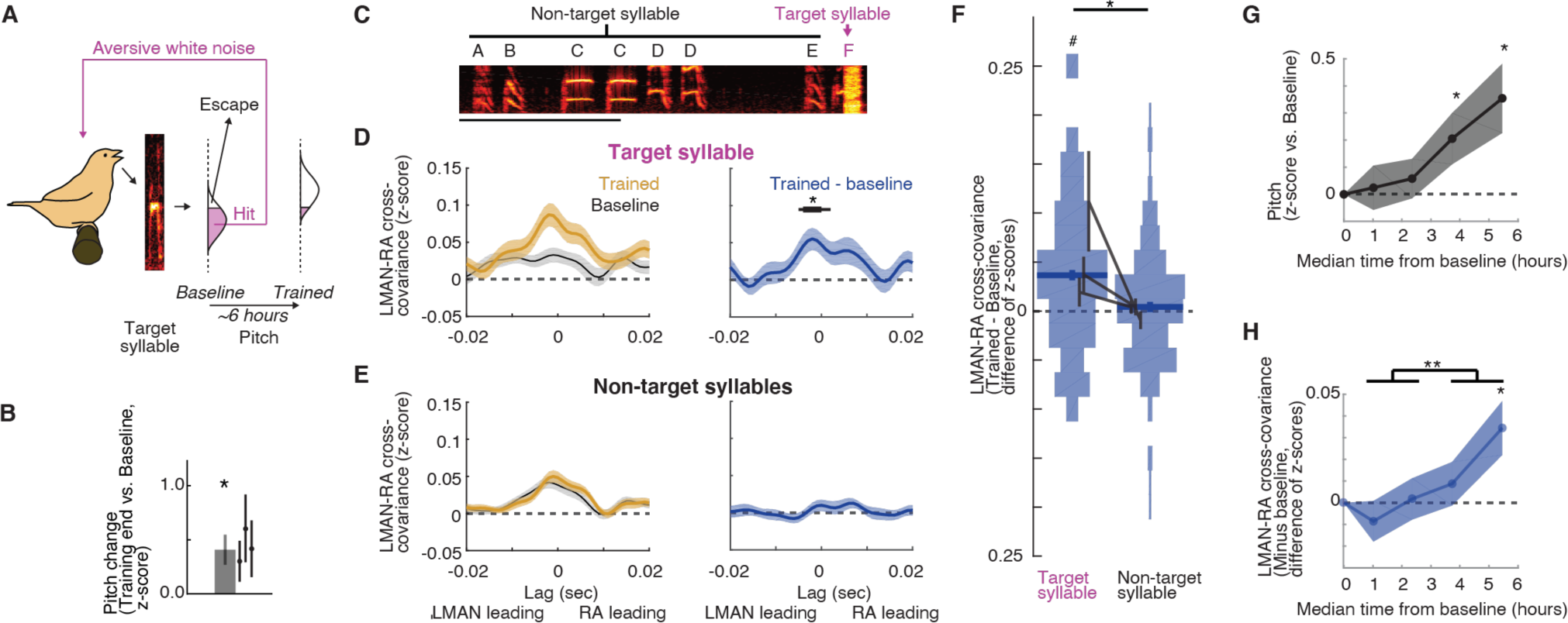
LMAN-RA co-fluctuations are enhanced during learning. (A) Pitch training paradigm. Aversive white noise feedback (“hit”) is delivered during renditions of a specific targeted syllable when it is either below or above a threshold (pink fill in histogram), depending on whether the objective is to train pitch shifts up or down, respectively. The schematic shows training for an upward pitch shift. Over the course of training (3-10 hours, mean ∼6 hours), birds progressively modify pitch in the direction that escapes WN (compare “Trained” vs. “Baseline” pitch distributions). (B) Summary of pitch change across experiments (mean +/S.E.M., N = 11 experimental trajectories over 3 birds); individual points represent the mean for individual birds. *, P<0.05, Wilcoxon signed-rank test. (C) Spectrogram of a song bout in an example experiment, with syllables labeled “Target” or “Non- target”, depending on whether they were targeted with WN. Scale bar, 500ms; y axis ranges from 0.5 to 7.0 kHz. (D) Change in cross-covariance during learning for targeted syllables. Left: mean +/- SEM cross- covariance for “Baseline” and “Trained” periods (last quarter of renditions during the training session) across experiments (N = 38 LMAN-RA site pairs, 11 experiments, 3 birds). Right: mean +/- SEM change in cross-covariance over the course of training. Black bar indicates time bins with values significantly different from zero (thin, p < 0.05; thick, p < 0.005). (E) Same as (D), but for non-targeted syllables. For a given experiment, there was only one targeted syllable, but multiple non-targeted syllables (mean, 4.2 syllables). Thus, for each LMAN-RA site pair, data were first averaged across all non-targeted syllables before plotting (N = 38 pairs, 11 experiments, 3 birds). (F) Summary of change in LMAN-RA cross-covariance during training across targeted and non- targeted syllables. For each combination of paired LMAN-RA sites and syllables, we computed the average change in cross-covariance in a 15 ms window centered at the peak of the average cross- covariance (-3 ms) at the end of training [N = 38 (Target) and 158 (Nontarget) combinations of paired sites and syllables, across 11 experiments in 3 birds]. *, p < 0.05, mixed effects model (fixed intercept and effect of syllable type; random effect of intercept and syllable type grouped by experiment ID). #, p < 0.05, mixed effects model (fixed intercept and random effect of intercept grouped by experiment ID). (G) Time course of pitch change. Each learning trajectory was divided into four bins (with equal number of renditions); average pitch change in each bin is plotted at the median time across trajectories (N = 10 experiments in 3 birds; this plot excludes one of the 11 experiments in panels B- F for which we failed to collect neural data throughout the entire training period). (H) Time course of change in LMAN-RA cross-covariance for the targeted syllable for the same experiments illustrated in (G) (N = 37 LMAN-RA site pairs, 10 experiments, 3 birds).

### The strength of LMAN-RA co-fluctuations and the strength of adaptive motor bias are associated on a rendition-by-rendition basis

If changes to LMAN-RA cross-covariance reflect top-down commands that adaptively bias RA activity and pitch during learning, then for each targeted syllable, rendition-by-rendition variation in the strength of this signal should be associated with rendition-by-rendition variation in the magnitude of pitch shifts in behavior. To test this prediction, we took advantage of the rendition-to-rendition variation in pitch shift that birds naturally exhibit (Fig. 3A, large spread of pitch values in both baseline and trained periods) to divide interleaved syllable renditions into two groups, based on whether the rendition’s pitch was shifted in the adaptive direction by more or less than the median amount. (Fig. 3A; “baseline”: using last half of renditions before training onset; “trained”: using last quarter of renditions during training period). We found that renditions with pitch biased more strongly in the adaptive direction were associated with greater changes to cross-covariance during learning (example experiment in Fig. 3A, B; summary in Fig. 3C), regardless of whether birds were trained to adaptively shift pitch up or down (Fig. S3). This finding that the strength of LMAN-RA cross-covariance and adaptive modifications to pitch co-evolve on a rendition-to-rendition basis suggests that LMAN-RA co-fluctuations reflect a rapidly varying topdown signal that adaptively biases motor performance during learning.

**Figure 3.**
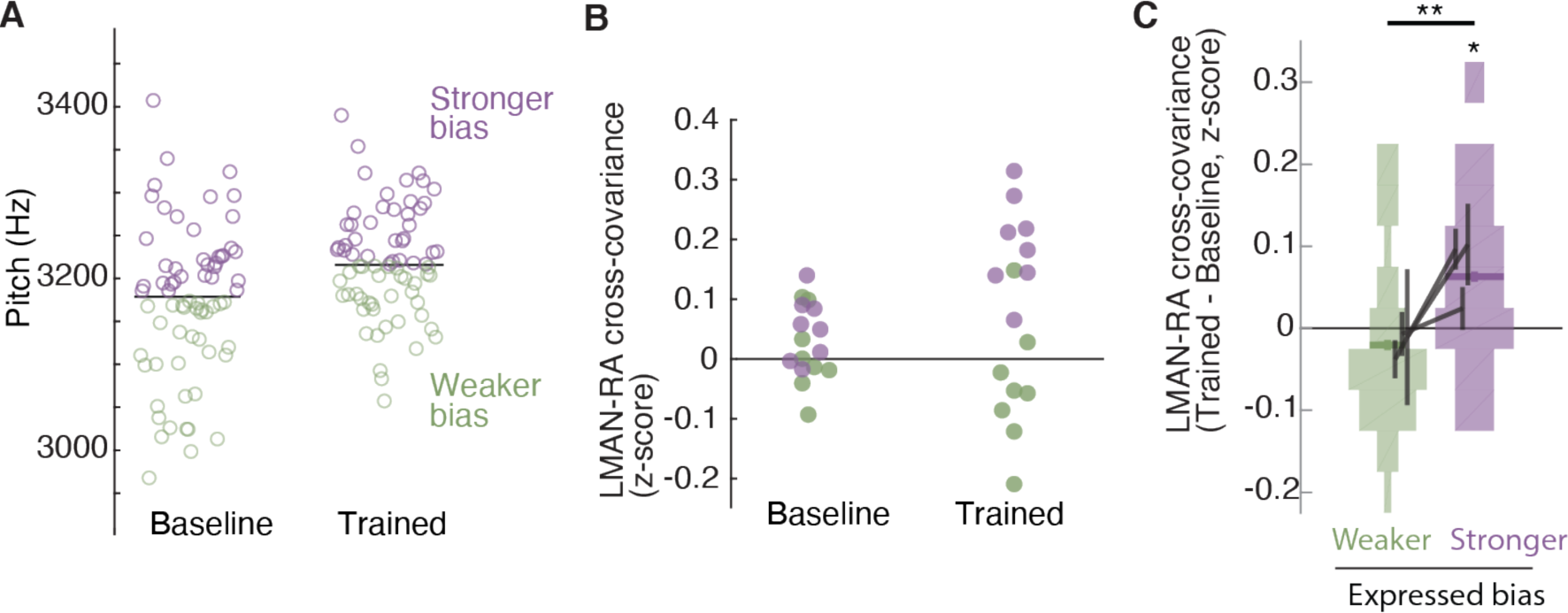
The strength of LMAN-RA co-fluctuations and the strength of adaptive motor bias are associated on a rendition-by-rendition basis. (A) Example experiment. Pitch of individual renditions during baseline and at the end of training in an experiment in which WN feedback targeted lower pitch renditions. Renditions are grouped into two sets, “Stronger bias” (purple) and “Weaker bias” (green) based on pitch deviation from the median pitch. Renditions in the “stronger” and “weaker” groups are interleaved in time, and taken from periods with stable pitch, so that any differences in LMAN-RA cross-covariance between these groups reflects rendition-by-rendition variation, not slower “drift” due to learning. We used the last half of baseline renditions and the last quarter of the training renditions. (B) Example experiment (same as in panel A), plotting LMAN-RA cross-covariance for individual pairs of LMAN-RA sites (filled circles). Each datapoint represents the mean cross-covariance for renditions with either Stronger bias (purple) or Weaker bias (green). Thus each site pair contributes two points each to the “Baseline” and “Trained” data (one purple, one green). The renditions used to calculate cross-covariance are the same as in panel A. (C) Summary of change in LMAN-RA cross-covariance during training (Trained - Baseline), measured separately for renditions expressing “stronger” or “weaker” bias (N = 38 LMAN-RA site pairs, 11 experiments, 3 birds). *, p < 0.05, LMAN-RA cross-covariance (Trained-Baseline) modeled with fixed intercept and random intercept grouped by experiment. **, p < 0.005, Stronger-Weaker modeled with fixed intercept and random intercept grouped by experiment ID.

### Learning-related increases in LMAN-RA co-fluctuations are context-specific

Prior studies revealed that pitch modifications for a given targeted syllable are dependent on sequential context, or the sequence of syllables in which the target syllable is embedded (Hoffmann and Sober, 2014; Tian and Brainard, 2017); this prompted us to examine whether increases in LMAN-RA cross-covariance are similarly context-specific. To do so, we took advantage of the fact that if aversive feedback targets a syllable (e.g., B) only when it is sung in the targeted context (e.g., AB) and not in other non-targeted contexts (e.g., XB, YB, etc), then the resulting pitch modification will be greatest when the target syllable is sung in the target context (Hoffmann and Sober, 2014; Tian and Brainard, 2017). In a subset of experiments, we therefore provided WN feedback in such a context-specific manner, with learning and non-learning contexts naturally interleaved from rendition to rendition (Fig. 4A, B). We found that average LMAN-RA cross- covariance increased over the course of training specifically for the targeted context, with no detected change for the same syllable in non-targeted contexts (Fig. 4C). Enhanced LMAN-RA cross- covariance is therefore flexibly linked to sequential contexts associated with learning.

**Figure 4.**
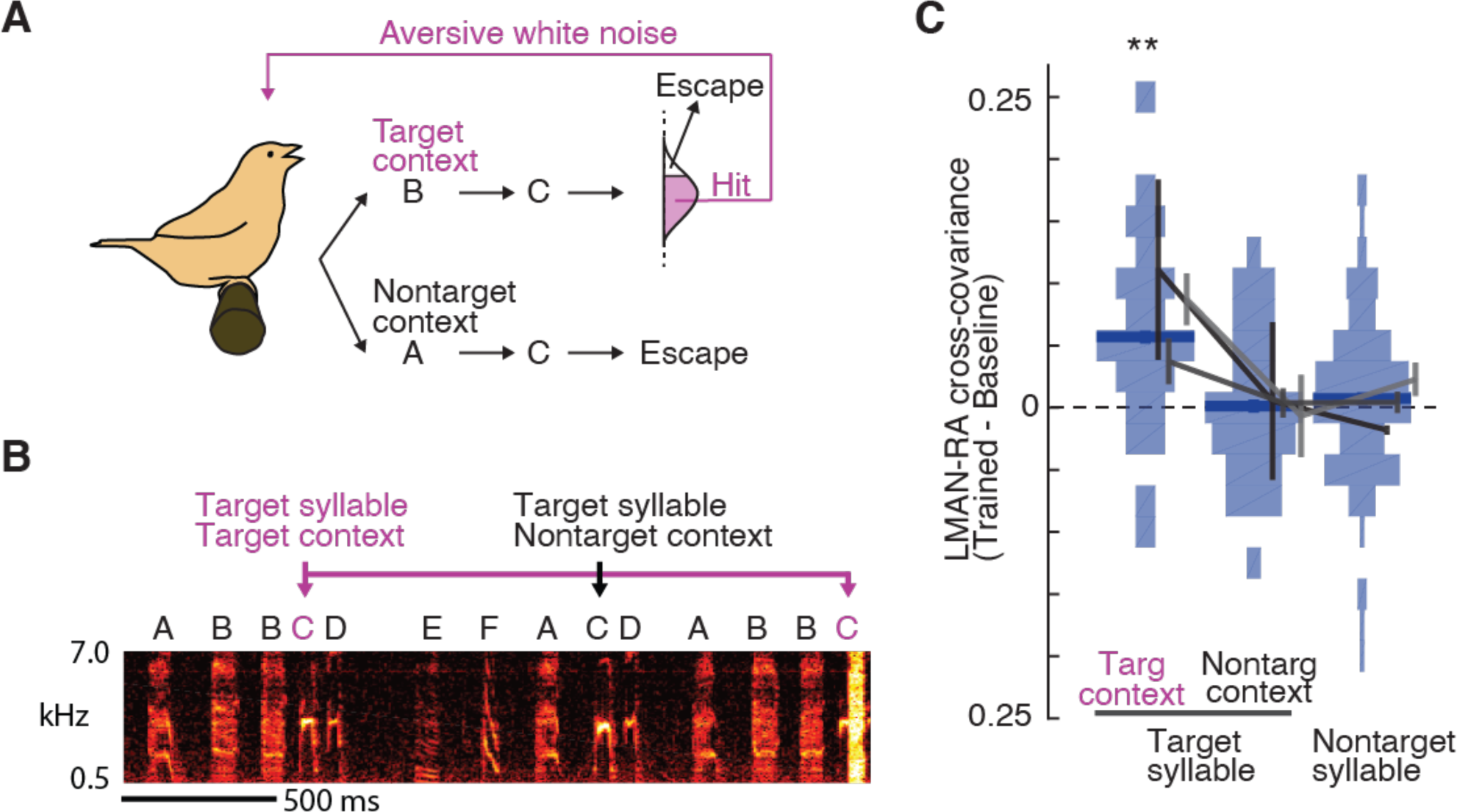
Learning-related increases in LMAN-RA co-fluctuations are context-specific. (A) Schematic of context-dependent training. In this example experiment, pitch-contingent WN (“hit”) was provided for renditions of the target syllable (C) only in the “Target” context (BC). WN was never provided when C was sung in “Nontarget” contexts (e.g., AC). (B) Spectrogram for the same example experiment as in panel A, illustrating WN delivered only for syllable C in the target context BC. (C) Summary of change in LMAN-RA cross-covariance during training (N = 24 LMAN-RA site pairs for targeted syllable, targeted context, 35 pairs for targeted syllable, non-targeted context, and 112 pairs for non-targeted syllables, 7 experiments, 3 birds). **, p<0.005, mixed effects model (fixed effect of intercept and syllable-type; random effect of intercept and syllable type grouped by experiment ID).

### Adaptive bias is eliminated by disrupting LMAN activity in a narrow premotor window

The close relationship between changes to LMAN-RA cross-covariance and behavior is consistent with a model in which LMAN provides a temporally localized top-down command that has a transient, causal and adaptive influence on the immediately following targeted syllable. However, prior studies manipulating LMAN activity to probe its contributions to pitch modification have used pharmacological manipulations that act on the timescale of minutes to hours (Amberg and Lindefors, 1989; Andalman and Fee, 2009; Charlesworth et al., 2012; Tian and Brainard, 2017; Warren et al., 2011), and thus lack the temporal resolution to assess the moment-by-moment relationship between neural activity and behavior. We reasoned that electrical microstimulation of LMAN could potentially enable a more temporally precise disruption of neural activity and test of LMAN’s causal influence on RA. Stimulation of LMAN (Giret et al., 2014; Kao et al., 2005; Kojima et al., 2018), and interconnected song system nuclei (Ashmore et al., 2005; Fee et al., 2004; Vu et al., 1994) during baseline singing perturbs song acoustic features in a manner that is rapid and transient (timescale of 10s of milliseconds). LMAN microstimulation—with intensity titrated to minimize overt behavioral effects during baseline singing (see Methods)—thus has the potential to disrupt neural activity that may convey a motor biasing signal during learning, revealing only pitch modifications that are encoded in the downstream motor pathway (Andalman and Fee, 2009; Warren et al., 2011).

To evaluate the specificity of the top-down bias reflected in increased LMAN-RA cross- covariance, we stimulated LMAN during the target syllable’s premotor window (60ms window, centered 50ms prior to syllable onset) during a randomly interleaved 50% of renditions (“Stim”) in 2-4 hour long blocks (Fig. 5A, B). Compared to prior studies using spatially localized, unilateral microstimulation of LMAN, our stimulation experiments were designed to cause more global, bilateral disruption of LMAN activity (see Methods). We measured the rendition-by-rendition effect of stimulation by comparing syllable pitch on “Stim” renditions to pitch on randomly interleaved control renditions (“Catch”) for which no stimulation was delivered. During baseline, before the onset of pitch-contingent WN training, stimulation caused modest changes to pitch that varied across experiments, resulting in only small changes to average pitch (“Baseline” in example experiment in Fig. 5C, D; summary in Fig. 5E; see also (Giret et al., 2014; Kao et al., 2005; Kojima et al., 2018)]. In contrast, during learning, stimulation caused pitch to revert systematically towards its baseline value (Fig. 5C-E, compare “Stim” vs. “Catch”). This occlusion of pitch modification was quantitatively indistinguishable from that caused by pharmacological inactivation of LMAN or blockade of LMAN’s synaptic input to RA as previously reported in the same species and training paradigm [(Warren et al., 2011) (compare Figs. 5E and F)]. Thus, our findings indicate that patterned LMAN activity is required for adaptive biasing of vocal output during learning.

**Figure 5.**
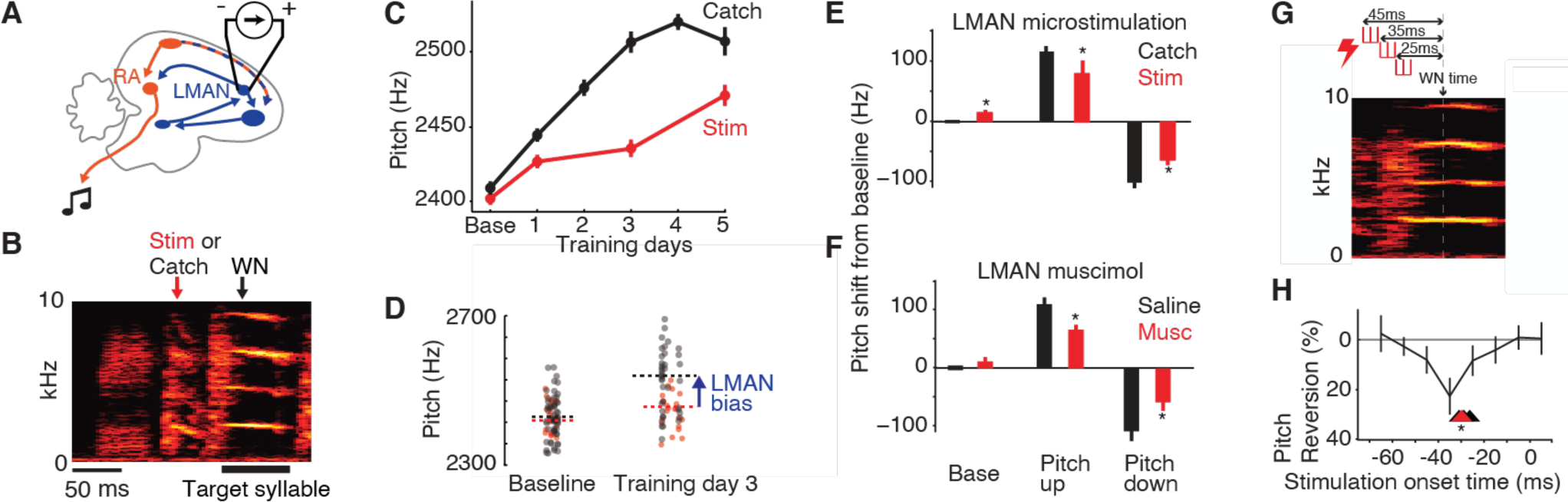
Adaptive bias is eliminated by disrupting LMAN activity in a narrow premotor window. (A) Schematic of electrical microstimulation in LMAN. Stimulation was used to disrupt LMAN activity at precise timepoints during singing. (B) Schematic of experimental design. To test for a causal contribution of LMAN premotor activity to pitch modifications during learning, on randomly interleaved renditions stimulation was either delivered [“Stim, during a 60ms window centered 50ms prior to syllable onset] or withheld (“Catch”). WN depicts average time of WN feedback for this experiment. (C) An example experiment in which pitch was driven up during training over five days, with LMAN microstimulation performed on the baseline day and training days 1, 3, and 5. (D) Scatterplot showing pitch of individual renditions from the experiment in panel C on randomly interleaved stimulated (red) and catch renditions (black) during ∼4hr periods at baseline and on the third day of training. The magnitude of LMAN bias is inferred as the extent to which learning is occluded due to stimulation. (E) Summary of effects of LMAN microstimulation on pitch across all experiments, plotted for baseline (“Base”), and end of training for experiments where we trained pitch up (“pitch up”) or down (“pitch down”). *, p<0.05, t-test comparing Catch to Stim. (F) Same as panel E, but for experiments in which LMAN was instead inactivated pharmacologically for a few hours using muscimol or lidocaine (both labeled “Musc”). Pharmacological perturbation had quantitatively similar effects on pitch compared to electrical microstimulation in panel E. Data were previously published (Warren et al., 2011). (G) Schematic of experiment using short-duration microstimulation (10 ms, red hash marks) applied at varying timepoints in the targeted syllable’s premotor window. Onset timing of stimulation was varied from -65 ms to +5ms relative to the expected timepoint of WN delivery (“WN time”). Three example microstimulation trains are shown, with onsets at -45 ms, -35 ms, and -25ms. (H) Summary of the effects of microstimulation on pitch as a function of stimulation onset time (mean pitch reversion +/- SEM). Stimulation trains that began 35 ms prior to FF measurement caused significant reversion of learning toward baseline. *, p<0.05. Stimulation trains slightly earlier and later also tended to cause a reversion of FF toward baseline, but did not achieve statistical significance. Arrowheads depict the minimum latencies across experiments (N=4) at which significant reversion occurred, including the experiment in panel G (red).

To precisely localize the temporal window in which LMAN contributes to adaptively biasing motor output, we systematically varied the timing of stimulation relative to the targeted syllable, applying short stimulation trains (10 ms) at times ranging from -65 to +5 ms relative to the time point at which pitch modifications in behavior are maximum [Fig. 5G,H, average time of WN delivery, labeled “WN time” (Charlesworth et al., 2011)]. Stimulation caused reversion of learning only if applied within a narrow premotor window immediately prior to this time point (Fig. 5H, - 50 to -20 ms relative to “WN time”, peaking at -35ms,). The minimum latency across experiments between stimulation and pitch reversion was ∼30 ms (Fig. 5H, range 26-31 ms across four experiments), matching prior estimates of the premotor delay between LMAN activity and song (Giret et al., 2014; Kao et al., 2005; Kojima et al., 2018). Combined with findings from pharmacological perturbations within this circuit and with our analysis of changes to LMAN-RA cross-covariance during learning, these microstimulation experiments indicate that LMAN exerts a moment-by-moment, top-down influence on RA to implement adaptive motor bias.

## Discussion

Using paired recordings from LMAN and RA in singing birds, we identified a neural signature of a top-down influence of LMAN on RA, quantified as a short latency, LMAN-leading peak in the cross-covariance of neural activity. This LMAN-RA cross-covariance peak is present even at baseline prior to learning (Fig. 1), consistent with an ongoing role for LMAN in driving renditionby-rendition exploratory variation in behavior (Kao et al., 2005; Olveczky et al., 2005). Strikingly, LMAN-RA cross-covariance strengthens in a premotor window closely linked to the individual movement (syllable, Fig. 2), rendition-by-rendition variation in the magnitude of adaptive pitch modifications (Fig. 3), and sequential context (Fig. 4) associated with learning. Moreover, temporally localized perturbation of LMAN activity specifically within this premotor window causes rapid and transient occlusion of learned changes to pitch (Fig. 5). Combined, these results indicate that LMAN enables learning by conveying a dynamic top-down command to RA that varies on the timescale of individual movements and is flexibly linked to contexts associated with learning.

### Adaptive bias through dynamic top-down influence on primary motor area output

Our finding that LMAN-RA cross-covariance is a temporally precise neural correlate of adaptive behavioral modifications extends previous work that identified enhanced inter-area cofluctuations during learning (Koralek et al., 2013, 2012; Lemke et al., 2019; Makino et al., 2017; Sawada et al., 2015; Veuthey et al., 2020; Wagner et al., 2019). In particular, prior studies left unresolved whether such co-fluctuation signals reflect the moment-by-moment implementation of adaptive changes to movements instead of other slower modulatory processes such as those related to motivation, vigor, attention, motor preparation, or other kinds of global brain state changes (Cowley et al., 2020; Hikosaka et al., 2006; Ignashchenkova et al., 2004; Jaffe and Brainard, 2020; Kawashima et al., 2016; Muller et al., 2005; Noudoost et al., 2010; Sawada et al., 2015; Stavisky et al., 2017). It is possible that inter-area co-fluctuations in other systems similarly reflect dynamic top-down shaping of primary motor output in driving behavioral adaptation. Moment-by-moment and context-specific links between Close links between measures of inter-area co-fluctuations and behavior, similar to the linkthose we found between LMAN-RA cross-covariance and behavior pitch shift during learning, may be detected most readily for forms of learning that involve modification of specific movement parameters with high temporal (Kawai et al., 2015; Medina et al., 2005, 2000; Narayanan and Laubach, 2006; Rueda-Orozco and Robbe, 2015) or contextual specificity (Howard et al., 2012; Rochet-Capellan et al., 2012; Rochet-Capellan and Ostry, 2011; Wainscott, 2004), similar to song learning (Charlesworth et al., 2011; Lipkind et al., 2017; Ravbar et al., 2012; Tchernichovski et al., 2001; Tian and Brainard, 2017; Tumer and Brainard, 2007) .

### Neural mechanisms underlying an adaptive top-down influence of LMAN on RA

Our finding of learning-related increases in LMAN-RA cross-covariance raises the question of what neural mechanisms could instantiate a changing top-down bias. A prominent hypothesis is that LMAN bias reflects learning mechanisms that reinforce LMAN activity patterns associated with “successful” behavioral variants (Andalman and Fee, 2009; Brainard and Doupe, 2000; Charlesworth et al., 2011, 2012; Doya and Sejnowski, 1998; Fee and Goldberg, 2011; Gadagkar et al., 2016; Kao et al., 2005; Kearney et al., 2019; Troyer and Bottjer, 2001; Troyer and Doupe, 2000a; Tumer and Brainard, 2007; Warren et al., 2011). Such reinforcement could lead to a variety of changes to LMAN activity patterns that could alter the influence of LMAN on RA. These possible changes include gross increases or decreases in firing rate across LMAN neurons, altered correlational structure across neurons in LMAN (Brown and Raman, 2018; Darshan et al., 2017; Kumar et al., 2010; Riehle, 1997; Woolley et al., 2014; Zandvakili and Kohn, 2015), or more complex or distributed changes to the temporal structure of LMAN neurons’ firing patterns (Kao et al., 2008, 2005; Kojima et al., 2013; Olveczky et al., 2005; Palmer et al., 2021). We did not detect any systematic learning-related increases or decreases in average LMAN firing rates during learning (Fig. S2), suggesting that any changes to LMAN activity during learning may be heterogeneous or distributed in a fashion that will require elucidation by future studies monitoring larger numbers of neurons during learning.

### A functional architecture with distinct substrates for fast and slow learning

Our finding of adaptive top-down commands in initial stages of learning suggests that motor skill learning is generally supported by a functional architecture in which motor biasing signals are generated by more frontal or executive circuits (analogous to the Anterior Forebrain Pathway of which LMAN is part) that are distinct from the circuits encoding well-learned movements. An advantageous feature of this hierarchical architecture may be to enable rapid behavioral modifications in early phases of learning via top-down signals that flexibly update in response to changing goals, contexts and reward contingencies, while allowing for the stable representation of “core” motor programs in primary motor areas (Tian and Brainard, 2017).

On the longer timescale over which expertise is developed, such top-down biasing may also supervise slower plasticity in primary motor areas to cause long-lasting updates to core motor programs. This plasticity mechanism may operate for both birdsong (Andalman and Fee, 2009; Fee and Goldberg, 2011; Moorman et al., 2021; Tian and Brainard, 2017; Troyer and Doupe, 2000b; Warren et al., 2011) and other sensory-motor skills (Kawai et al., 2015; Knudsen, 1994; Turner and Desmurget, 2010), and may underlie the observation that early and late stages of learning are often supported by different neural substrates for birdsong (Andalman and Fee, 2009; Aronov et al., 2008; Bottjer et al., 1984; Nordeen and Nordeen, 2010; Scharff and Nottebohm, 1991; Warren et al., 2011) and other sensory-motor skills (Dayan and Cohen, 2011; Doyon and Ungerleider, 2002; Hikosaka et al., 2002; Kleim, 2004; Shadmehr and Holcomb, 1997).

## Methods

### Animal subjects

We used adult male Bengalese finches (*Lonchura striata domestica, N* = 8) that were bred in our colony and housed with their parents until at least 60 days of age. During experiments, birds were housed individually in sound-attenuating chambers (Acoustic Systems) on a 14h/10h light/dark cycle with food and water provided ad libitum. All experiments were performed on “undirected” song (i.e., with no female present). All procedures were in accordance with protocols approved by the University of California, San Francisco Institutional Animal Care and Use Committee.

### Pitch training paradigm using closed-loop reinforcement

We used a modified version of EvTaf (Charlesworth et al., 2012, 2011; Tian and Brainard, 2017; Tumer and Brainard, 2007; Warren et al., 2011), a custom-written Labview program (National Instruments), to monitor song and deliver white noise feedback in a closed-loop fashion during training (Ali et al., 2013; Andalman and Fee, 2009; Charlesworth et al., 2012, 2011; Tian and Brainard, 2017; Tumer and Brainard, 2007; Warren et al., 2011). Briefly, song was recorded with an omnidirectional lavalier microphone (Countryman), bandpass filtered between 75 Hz and 10 kHz, and digitized at 32 kHz. To detect a specific segment of song (i.e., within a targeted syllable) for targeted reinforcement, the spectrum of each successive 8ms segment of ongoing song was tested online for a match to a spectral template constructed to discriminate the targeted segment from all other song segments. Successful match was based on threshold crossing of the Euclidean distance between this song segment and template. A match signaled detection of the targeted syllable. Then, fundamental frequency (FF), or pitch, of the matching segment was compared to a preset FF threshold. If FF was below threshold (in experiments driving upwards shifts in FF), or above threshold (in experiments driving downwards shifts in FF), white noise feedback [WN, 40-60 ms at 90-95 dB(A)] was delivered with <1 ms latency. Feedback renditions were termed “hits.” Threshold was chosen to maintain ∼70% hit rate.

For context-dependent training, this paradigm was modified so that pitch-contingent WN was delivered only if the Target syllable was sung in a specific Target context, defined by the identity of the sequence of syllables directly preceding the Target syllable. Sequential context was defined for each rendition of the Target syllable and detected online by extending the spectral template matching algorithm (above) to detect conjunctions of the current and preceding syllables. These methods were the same as those used in (Tian and Brainard, 2017).

All training trajectories for neural recordings were performed within a single day of continuous recording of neural and singing data. WN was turned on after sufficient baseline data were collected (15 to 40 song bouts). The average training duration was 6.4 hours (min, 3.0; max, 10.0) starting from when WN was turned on.

For LMAN microstimulation experiments, training duration was extended to up to four days with the goal of eliciting large shifts in pitch that would allow more robust measurement of behavioral effects of stimulation. This approach was chosen to be the same as in previous pharmacology experiments (Warren et al., 2011) which allows direct comparison of behavioral effects across these different manipulations.

For all pitch training experiments, we initially set the WN pitch threshold to a hit rate of ∼70%. Successful learning leads to a progressive decrease in the hit rate; therefore, the pitch threshold was updated over the course of training to maintain a hit rate of 70%.

### Offline analysis of song

For analysis of song recorded simultaneously with neural recordings, we used song acoustic data recorded using the same Intan acquisition system used for collecting neural data (see below), to ensure temporal alignment of neural and singing data. Audio signals were acquired with an electret microphone (CUI), amplified (MAX4466, Adafruit), and digitized at 30 kHz. For analysis of song during LMAN stimulation, we analyzed data saved using the Labview training program described above.

Syllable pitch was calculated in the following manner (Charlesworth et al., 2011). For each syllable rendition, we calculated a spectrogram using a Gaussian-windowed (σ = 1 ms) short-time Fourier transform (window size = 1024 samples; overlap = 1020 samples; sampling rate = 32 kHz). Within each time bin, FF was defined as the frequency corresponding to peak power of the first harmonic, estimated using parabolic interpolation. FF for the rendition was then calculated as the mean FF across time bins for a fixed window defined relative to syllable onset. We excluded from analysis the two syllables directly following the target syllable, to avoid potential acute effects that may occur due to WN at the target syllable (Sakata and Brainard, 2008, 2006). We similarly excluded introductory notes and call-like syllables, which both consist largely of broadband noise and lack well-defined pitch.

### Electrode array microdrives for neural recordings

Custom microdrives, inspired by (Vandecasteele et al., 2012), were constructed to hold a custom-built array of tungsten electrodes. For one bird we used two 0.5 MOhm and two 6.0 MOhm electrodes in each array organized in a diamond formation (see below for spatial spread).

For other birds we used four (diamond) or five (pentagon) 0.5 MOhm electrodes. In all cases we used tungsten electrodes from Microprobes (WE30010.5F for 0.5 MOhm, and WE30016.0F for 6.0 MOhm). Microdrives consisted of a movable shuttle onto which electrodes were fixed, allowing manual adjustment of the position of electrodes along the z-axis with a resolution of 26.5 µm (one-eighth turn of a 00-120 screw). Electrodes were stabilized in the horizontal (x-y) plane by passing them through tight-fitting polyimide tubes (0.0056” ID, 0.00075” WD) glued to the static parts of the drive. Silver wires (diameter, 0.003” bare, 0.0055” teflon-coated) connected each electrode to a separate pin on an Omnetics connector (A79042-001). Low impedance reference electrodes were made by cutting tungsten electrodes described above to a blunt tip. Electrodes for the two microdrives (LMAN and RA) were wired to different channels on the same connector.

Electrodes were positioned in the array such that their tips were in the same horizontal plane. Electrodes were positioned so that there was greater spread along the anterior-posterior axis (0.4 – 0.5 mm) than the medial-lateral axis (0.2 – 0.3 mm) to account for the expectation of greater variability in targeting along the anterior-posterior axis, since position in this axis was further from stereotaxic zero than position in the medial-lateral axis (see below for coordinates).

### Implantation of recording microdrives

Implants were performed in the left hemisphere in the following order within a single surgical session with birds under anesthesia: RA microdrive, LMAN microdrive, reference electrode, shared connector. All wiring was completed before implantation. The location of RA was confirmed by electrophysiological targeting. Single carbon-fiber electrodes (Kation Scientific, Carbostar-1) were lowered gradually at candidate x-y locations to depths where RA was expected. RA was detected based on the presence of its characteristic tonic spiking activity (10- 20 Hz). The x-y location of the center of RA was determined as where the extent of tonic activity extended over a depth of at least 400 µm; we performed up to three penetrations at different x-y locations to determine the best estimate of RA’s center, which we found to be 2.05-2.15 mm lateral, 0.04 mm posterior to Y0 (i.e., caudal point of the intersection of the midsagittal and transverse sinuses), 2.75-3.15 mm ventral to the brain surface, with beak angle at 42 ° [by definition, vertical (beak pointing down) was 0° and horizontal was 90°].

A microdrive was then implanted with electrode tips 1.2 mm above the dorsal edge of RA. Next, a microdrive was implanted over LMAN using stereotaxic coordinates: 1.37 mm lateral, 5.18-5.27 mm anterior, 1.25 mm ventral, at a 50° beak angle. This depth was expected to place the electrode tips ∼0.5 mm above LMAN. The reference electrode was implanted directly underneath the skull but with minimal penetration of brain, either over the cerebellum or directly between the LMAN and RA implants. The connector was fixed to the skull over the right hemisphere. Dental cement [Coltene Hygenic] was used to secure all implants to the skull; small holes were made in the upper layer of the skull into which dental cement could flow before curing to increase implant stability. A plastic tube, glued to the dental cement base of the implant and surrounding the implant, was used as a protective cap.

### Electrophysiological recordings

After subjects recovered and were singing (1-2 days), they were tethered and then handled a few times a day for acclimatization. About a week or two after surgery birds consistently sang within tens of minutes after being handled, allowing recording sessions to begin. On each recording session, starting immediately after lights being turned on in the morning, we slowly lowered (<20 µm/sec) the electrodes from their resting positions towards LMAN and RA. Localization within these nuclei was assessed by evaluating tonic activity (RA), song-locked firing rate modulation (LMAN and RA), and depth. Post-hoc histological verification confirmed that recordings were within LMAN and RA (see below). At the end of the session, electrodes were raised to a position with the tip at >300 µm above the dorsal edges of RA and LMAN to minimize potential tissue damage within those areas. Recording sessions were separated by multiple days, and different depths were targeted.

Voltage signals were measured using a homemade lightweight headstage (Intan RHD2132 amplifier chip) and the Intan RHD2000 Amplifier Evaluation System. Signals were amplified, filtered (1Hz to 12000Hz pass band), and multiplexed on the headstage, then stored on hard disk for offline analysis.

A total of seven birds received dual LMAN and RA implants. For three of these birds, we were unable to obtain any appropriate data due to song deterioration after surgery (likely due to damage of HVC, RA or the HVC-RA tract). We obtained LMAN and RA recordings from the remaining four birds. For one of these birds we were only able to collect neural data during baseline singing, as signal strength had degraded, to the point where spikes were not detectable, before the rate of songs per day had recovered to a level sufficient for training pitch modification. Therefore, analyses of structure of LMAN and RA activity during baseline singing included all LMAN and RA recordings (including those not recorded concurrently) resulting in a total of N=30 LMAN and 52 RA multi-unit sites across four birds (ranging from 3-10 LMAN and 5-20 RA sites per bird). Analysis of LMAN-RA cross-covariance in learning experiments included only concurrent LMAN and RA recordings collected over a day of learning, resulting in a total of N = 38 pairs of LMAN and RA sites across 11 learning sessions in 3 birds. In each training session we recorded 1-2 sites in LMAN and 1-5 sites in RA, and considered all pairs of these LMAN-RA sites for LMAN-RA cross-covariance.

### Spike detection

Spikes were detected using the spike clustering software Waveclus (Chaure et al., 2018) run on MATLAB. Briefly, we detected putative spikes by amplitude threshold crossing (threshold of 2.5-3.5 x SNR, minimum refractory period 0.2 ms), mapped those spikes onto a feature space defined by wavelet coefficients, and then clustered spikes in this feature space. Because we did not observe spikes clustering into distinct classes, as we expected given our choice to use low- impedance electrodes, we pooled all spikes for a given channel in a given session into one multiunit cluster.

Prior literature suggests that a substantial portion of the spikes we recorded were from excitatory projection neurons. First, for LMAN, the ratio of projection neurons to interneurons has been estimated at around 3:1 (Bottjer et al., 1998) to 15:1 (Livingston and Mooney, 1997). Further evidence that excitatory neurons are more easily detected than inhibitory ones comes from reports of difficulty finding interneurons in slice preparations (Boettiger and Doupe, 1998; Livingston and Mooney, 1997). One study reported that LMAN projection neurons identified as projecting to RA (based on antidromic stimulation) have similar spiking activity to LMAN neurons whose projection status was not verified with antidromic stimulation, suggesting that extracellular recordings in LMAN tend to be dominated by projection neuron activity (Olveczky et al., 2005). For RA, the tonic firing we found in the multi-unit activity is characteristic of projection neurons (Leonardo and Fee, 2005; Sober et al., 2008; Spiro et al., 1999). Moreover, other studies isolating single neurons have reported low probability of finding putative interneurons in RA (Leonardo and Fee, 2005; Sober et al., 2008); indeed, in slice preparations the ratio of projection neuron to interneurons in RA was found to be around 30:1 (Spiro et al., 1999).

### Analysis of temporal structure of LMAN and RA activity during singing

To assess the temporal structure of activity separately in LMAN and RA, we computed the average firing patterns aligned to song motifs (stereotyped sequences of syllables) in Fig. 2B. Motifs were identified by visual inspection of a large corpus of song bouts for each bird (N = 8 motifs across 4 birds). For a given motif, the timing of syllable onsets and offsets can vary slightly across renditions and recording sessions. To account for this variation and temporally align activity across renditions of a given motif, we linearly time warped all renditions to an “reference” motif rendition, constructed with syllables and gaps matching the median value of syllables and gaps for that specific motif across all of its renditions. This alignment was performed by shifting each spike’s time so that its fractional time within its containing segment (i.e., syllable or gap) remained unchanged after time warping. See example spike trains preand post-warping in Fig. S1A.

To compute a smoothed firing rate function, activity on each rendition was smoothed by first binning spikes (1 ms) then convolving with a gaussian kernel (5ms S.D.). Rendition-averaged activity was then z-scored to facilitate comparison across recording sites, which may have different firing rates, relative to the mean and S.D. over the entire motif.

To assess the similarity of LMAN and RA activity patterns, we combined all recordings across sessions into separate LMAN and RA datasets, one for each motif, where activity was time warped, smoothed, and averaged over all sites (without first z-scoring) and then meansubtracted. The cross-correlation of the resulting mean LMAN and RA activity traces was computed using the matlab function XCORR, with argument scaleopt=’coeff’ to return as correlation coefficients.

For analysis of average LMAN and RA activity during baseline singing, to assess whether the average time lag of maximum LMAN-RA cross-correlation, across motifs, was significantly different from zero, we performed a Monte Carlo permutation test (Fig. S1F). We computed a null distribution under the null hypothesis that LMAN and RA activity patterns have the same temporal profile relative to song. On each shuffle iteration we randomized the assignment of rendition-averaged neural activity to brain region. We first generated a dataset consisting of average activity patterns, one for each combination of motif and brain area. On each shuffle iteration, we randomly reassigned the brain area labels; this was done independently for each motif, which ensures that the number of LMAN and RA sites assigned to each motif (and therefore bird) remained unchanged.. LMAN-RA cross-correlations were computed with this shuffled dataset for 10,000 random permutations. The probability of finding in the shuffled dataset an absolute time lag equal to or greater than the absolute time lag in the real dataset was taken as the two-sided p-value.

### Normalized cross-covariance

Normalized cross-covariance was computed for each pair of LMAN and RA sites for each syllable’s premotor activity (neural activity extracted from 100 ms to 0 ms preceding syllable onset). This calculation was done separately for baseline and training. For baseline, we used the last half of renditions, to minimize potential drift in recordings from lowering electrodes into position at the start of the session. For training, we used the last quarter of the renditions to take the window of maximal learning.

Normalized cross-covariance was calculated in a similar manner to previous birdsong studies (Hahnloser et al., 2006; Kimpo et al., 2003). We first calculated the average cross- correlation:

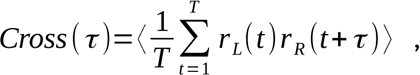

where *r _L_*(*t* ) *,r _R_* (*t* ) are binned spike counts at time bin *t* for LMAN and RA respectively, *T* is number of time bins (in a single rendition), τ is lag between LMAN and RA data in units of time bins, and ⟨ ⟩ indicates average over all renditions. We used time bins of 2.5 ms.

The cross-covariance was computed to estimate the extent to which deviations of firing rates in LMAN and RA from their respective means are associated:

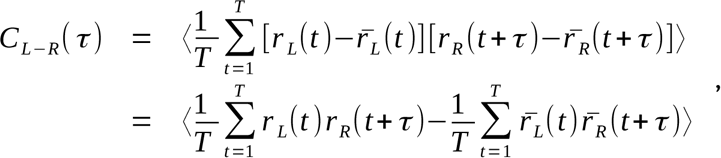

Where *r̄*_*L*_(*t* ) *r̄*_*R*_(*t* ) are rendition-averaged firing rates in LMAN and RA.

The second term on the right hand side 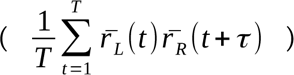 is the cross-correlation of the average firing rates, since it measures the average similarity of LMAN and RA activity while removing the contribution of shared, within-rendition variation in LMAN and RA. It can be estimated by calculating a shuffled cross-correlation (Brody, 1999; Hahnloser et al., 2006; Kimpo et al., 2003; Perkel et al., 1967). The shuffled cross-correlation (or “shift predictor”) is performed using data shuffled such that rendition *n* for one recording site (either LMAN or RA) is compared to a temporally adjacent rendition (i.e., *n+1* or *n-1*) for the other site:

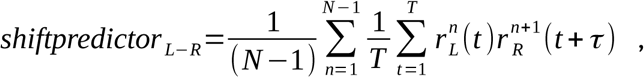

where *n*=1… *N* indexes the rendition, and the *L*−*R* subscript indicates that LMAN renditions (*n*) are chosen to be the one directly preceding the RA rendition ( *n+1*). We also computed the *R*− *L* shift-predictor:

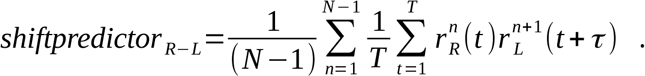

The final shift predictor was the average of these two:

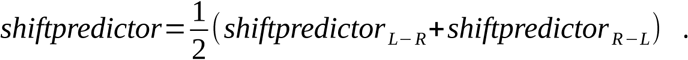

Subtracting the shift predictor from the average cross-correlation gives the cross-covariance:

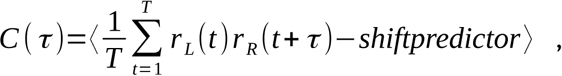

in units of spikes^2^. To rescale cross-covariance in units more easily comparable across the dataset we normalized cross-covariance relative to the standard deviation of the cross-covariance across all the datapoints (i.e., each combination of rendition *n* vs. rendition *n+1*) in the shuffled dataset. This effectively z-scores the cross-covariance relative to the mean and standard deviation of the shuffled distribution:

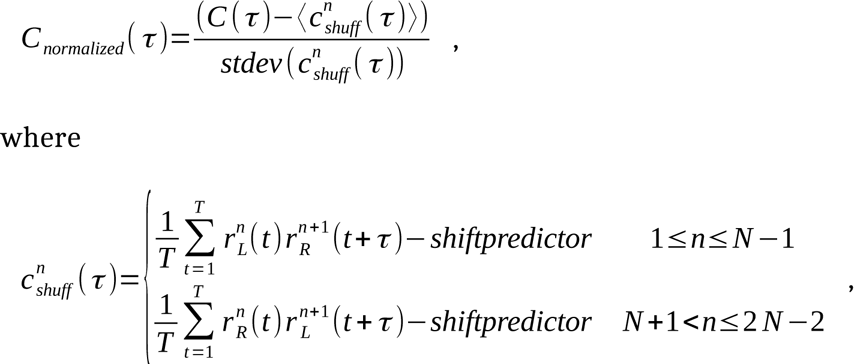

and *stdev* is the standard deviation over all *n* (from *0* to *2N-2*), for the time bin τ for the shuffled data:

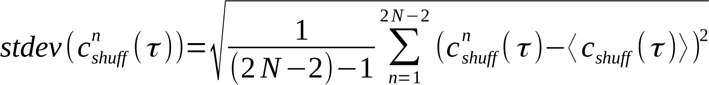

Cross-correlation functions were first linearly interpolated to 1 ms resolution then smoothed with a gaussian kernel (S.D. = 5 ms) before converting to normalized cross-covariance. Calculation of cross-correlations were implemented using the MATLAB function *XCORR* (with *SCALEOPT*=’unbiased’). Normalized cross-covariance was performed separately in 60 ms windows, sliding over 5 ms timesteps, in each syllable’s premotor window (such that the earliest window spanned from 100 ms to 40 ms preceding syllable onset, and the latest window from 60 ms to 0 ms). The cross-covariance functions over all 60ms windows were then averaged to generate one cross-covariance function for a given 100ms premotor window.

To summarize the strength of the LMAN-leading peak in a scalar value, we took the average within a 15 ms window centered at the time lag of peak normalized cross-covariance at baseline across all syllables (3 ms, LMAN leading).

For the analysis splitting interleaved renditions by pitch into two groups (“Stronger” and “Weaker” bias), we split renditions by comparing their pitch to the median pitch. If pitch deviated from the median pitch in the adaptive direction (that is, escaping WN), then the rendition was considered to express stronger behavioral bias; if pitch was in the other direction, then weaker bias. This grouping was performed separately for baseline and WN renditions relative to their respective median pitch values, ensuring that the resulting groups consisted of interleaved renditions. As in the regular analysis of learning, we used the last half of the baseline renditions and the last quarter of the training renditions. For the “Baseline” dataset, the adaptive direction was set the same as that for the “Trained ”dataset so that we could specifically measure learning- related change in the relationship between pitch and neural activity by subtracting baseline from training measurements. Normalized cross-covariance was computed as before, separately for each dataset group. Shuffled renditions were constrained to be only those pairs of renditions overlapping with the renditions included in a given group. For example, if rendition *m* for LMAN is included in the group, then the corresponding shuffle renditions will be *m* vs *m-1* (LMAN vs RA) and *m* vs *m+1*. At dataset edges, when *m=1*, rendition *N* was used for RA, and when *m=N*, rendition 1 was used.

### Electrical microstimulation of LMAN

We used a modified version of EvTaf that enabled electrical stimulation to be delivered independently of white noise feedback. In order to stimulate at a controlled time relative to the syllable targeted for learning, we detected “predictor” syllables that consistently preceded the targeted syllable. We then delivered stimulation trains (parameters described below) at a fixed delay from this detection so that stimulation began at a premotor latency prior to the WN trigger time. Stimulation renditions were randomly interleaved with catch renditions in which no stimulation was delivered. We stimulated a randomly interleaved 50% of renditions of the targeted syllable only during specific two-to-four hour intervals on specific days at baseline and after at least two days of training.

Custom-made 4-wire Pt/Ir microwire arrays (Microprobe; 25 micron diameter, 400-800 kOhm impedance wires) were surgically implanted bilaterally. The arrays were targeted using stereotaxic coordinates for the center of LMAN. Wires were electrically connected to a male Omnetics connector (A8391-001) that enabled electrical connection to an external lead. The four wires in each array were arranged in a rectangular pattern (250-500 micron separation on the rostro-caudal axis, 250-500 micron separation on the medial-lateral axis) for all but one bird.

Wire pairs at the same rostral/caudal or medial/lateral level were separated in depth by 0-250 microns. In one bird, the four wires were laid out in a linear array in which wires were separated by 250 microns along the medio-lateral axis, and neighboring wires were separated by 250 microns in depth.

After recovery from surgery, microstimulation trains (biphasic pulses, total biphasic pulse duration of 0.4ms, 200Hz frequency, 30-100 µA) were delivered to LMAN bilaterally by two separate microstimulators (A-M systems Model 2100). This bilateral stimulation contrasted to prior studies which showied that LMAN stimulation can induce large, rapid deflections in pitch (Kao et al., 2005) in three ways that were expected to enhance our ability to globally disrupt LMAN activity without inducing novel motor-driving patterns. First, the bilateral nature of our stimulation was expected to induce more global disruption compare to unilateral stimulation in (Kao et al., 2005). Second, we passed current between pairs of identical wires placed at different locations within LMAN (distance between electrodes >= 350 microns), rather than from a single electrode to ground; we expected our approach to perturb a larger volume of LMAN along the current path between the two electrodes. We used either a rectangular configuration, in which current was passed across a diagonal wire pair, or a linear configuration, in which current was passed across the two inner wires. We selected pairs of arrays for stimulation which elecitied minimal baseline pitch deviations. Third, we set the amplitude of stimulation current at a level below that which song stoppages or degradation of syllable structure occurred.

### Post-mortem localization of recording and stimulation sites

For both recording and microstimulation experiments, we marked the location of electrodes by first lesioning brain tissue and then performing histology to map those lesions relative to sites of recording (LMAN and RA) or microstimulation (LMAN). Lesions were performed by passing 100 uA current for four seconds. After lesions, birds were deeply anesthetized and perfused with 4% formaldehyde. Brains were removed and post-fixed for a few hours to overnight. We performed histology on sectioned tissue (40 µm thick, coronal). Electrode tips were localized by identifying lesions and tracts by identifying tissue damage. LMAN and RA were visualized by immunostaining for calcitonin gene related peptide (Sigma, RRID: AB_259091, 1:5000 to 1:10000) (Bottjer et al., 1997). For microstimulation experiments, we confirmed that lesions were in LMAN. For neural recording experiments, two lesion sites were made, one immediately dorsal and another immediately ventral to LMAN and RA, in order to retain the integrity of tissue within each area for histology. We confirmed in histology that lesions were indeed positioned dorsal and ventral to LMAN and RA, such that electrodes would be expected to be within these regions when at stereotaxic depths used during recordings.

### Statistical tests

The main recording results were analyzed using mixed effects modeling to capture potential heirarchical effects based on experimental session, because a given experimental session may contribute multiple pairs of sites, which are not completely independent. In general, we modeled responses (changes in LMAN-RA cross-covariance) with fixed effects for the covariate of interest, with random effects for intercept and the covariate of interest grouped by experiment ID. We used the Matlab function *FITLME* (with ‘STARTMETHOD’=’random’).

---

**Supplemental Figure 1.**
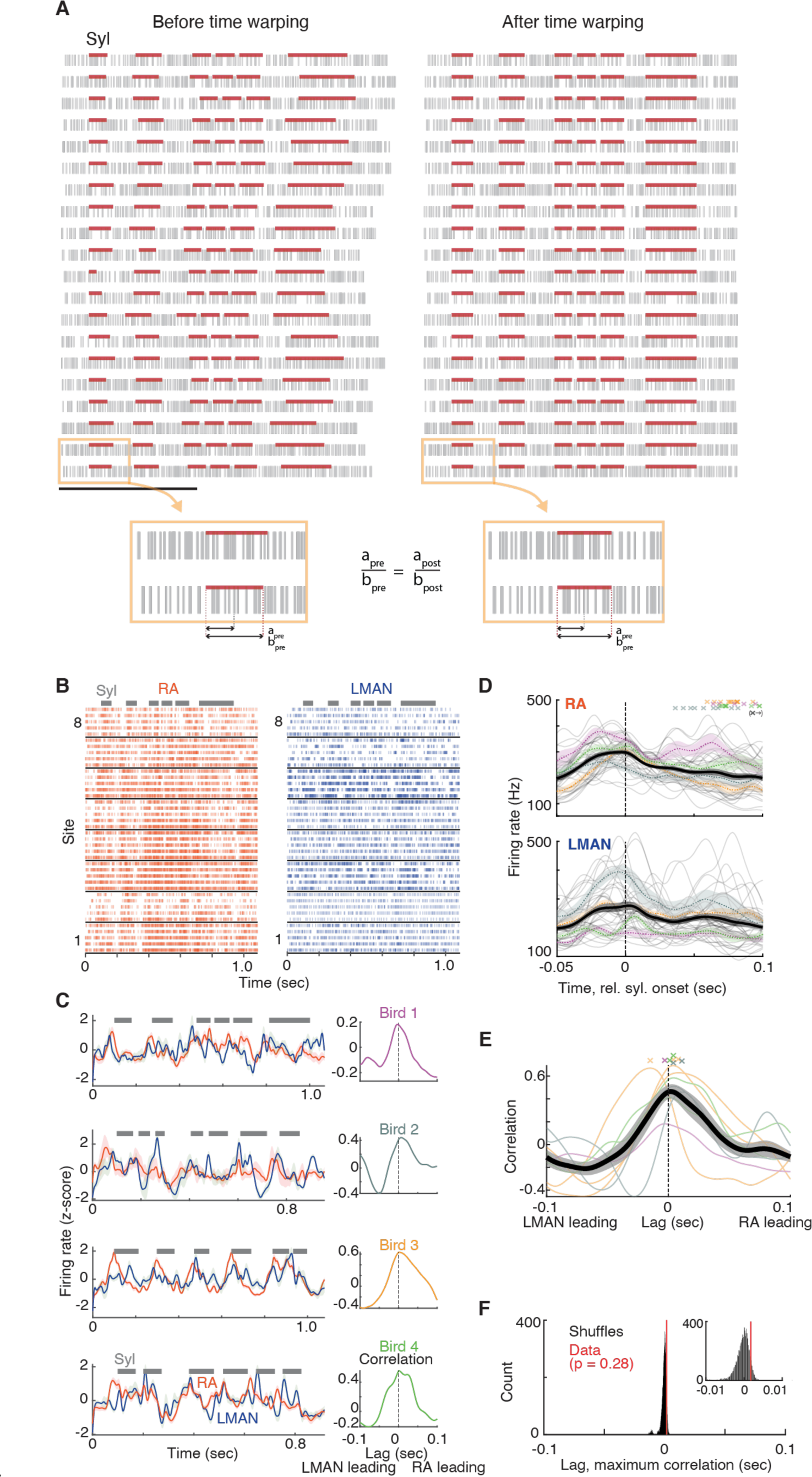
LMAN and RA average activity patterns align to each other with time lag close to zero. (A) Linear time warping methodology. Linear time warping was used to align all renditions of a given motif (across all renditions for all LMAN and RA recording sites) to a single reference rendition, before performing the analyses of temporal structure in this figure. Raster of spike times before (left) and after (right) time warping, for twenty example renditions (rows) of an example motif. Horizontal red bars represent times of syllables in this motif. Time warping aligns the spike times for all renditions to a common a median “reference” rendition of this motif (see Methods). The bottom two panels illustrates in detail that this time-warping procedures shifts spikes under the constraint that the spike’s fractional time within its containing segment (either syllable or gap) remains unchanged after shifting (a_pre_/b_pre_ = a_post_/b_post_). (B) Raster plots of spiking activity for eight (rows) example RA (left) and LMAN (right) sites aligned to a single motif, with 5 renditions shown for each site. Syllables are represented by black horizontal bars. (C) Left column, mean LMAN (blue) and RA (red) activity over all recorded sites, for four example song motifs, each from a different bird (rows). Syllable are indicated by grey bars. Right column, cross-correlation between mean LMAN and RA firing rate traces for the data on the left. N = 8 LMAN sites and 12 RA sites (for the motif for Bird 1), 3 LMAN and 5 RA (Bird 2), 9 LMAN and 15 RA (Bird 3), 10 LMAN and 20 RA (Bird 4). (D) Mean activity aligned to syllable onsets for all RA (top) and LMAN (bottom) sites. Thick, dark trace is mean +/- SEM activity taken over all syllables [N=39 (range 6-14 across birds), each represented as a thin gray trace showing mean across sites], extracted from all motifs (N=8 motifs for 4 birds, range 1-3 per bird). Colored, dashed traces show mean for each bird. Crossmarks on top panel represent offset of each syllable. Arrows indicate that offsets for four syllables are off the graph, at timepoints 0.11, 0.12 (orange), 0.11 (blue), and 0.19 (purple) seconds]. (E) Cross-correlation functions for all song motifs. Thin traces represent cross-correlations for each motif, colored by bird (same as panel B). Thick trace represents mean +/- SEM across motifs (N = 8 motifs). Crosses on top represent times of maximum correlation for each motif. N = 30 LMAN, 52 RA sites total (range 3-10 LMAN, 5-20 RA across 4 birds). (F) Comparison of the mean time lag of maximum correlation (red line, mean of cross marks in panel D) to a null distribution of time lags, representing the null hypothesis that there is no consistent time lag between LMAN and RA average activity patterns, estimated by recomputing (10000 times) the mean time lag of maximum correlation after shuffling the brain area “labels” associated with each rendition-averaged firing rate trace (see Methods).

**Supplemental Figure 2.**
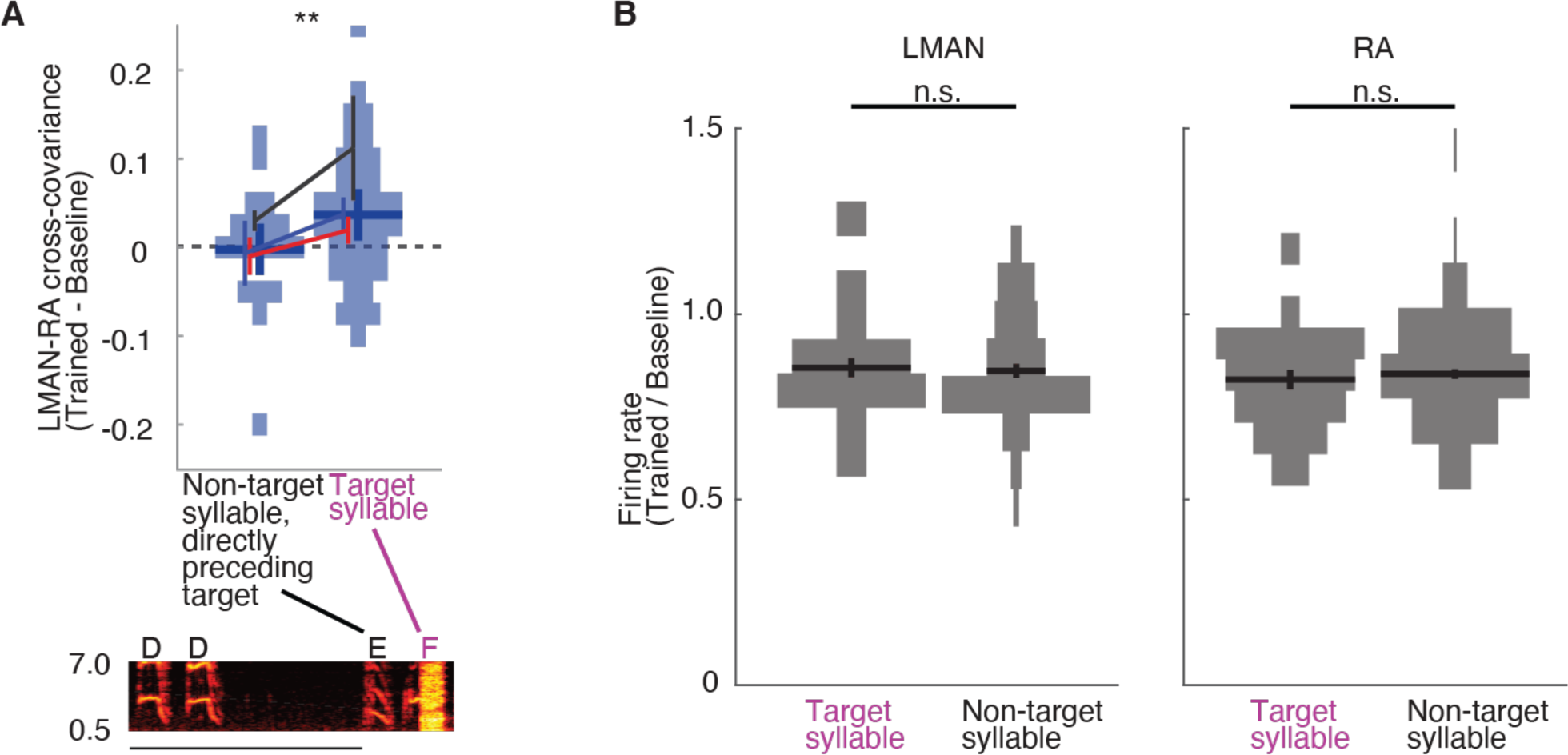
Specificity of learning-related change to LMAN-RA co-fluctuations (Related to Figure 2). (A) Change in LMAN-RA cross-covariance did not occur for non-targeted syllables sung directly preceding the targeted syllables in the song sequence. N=19 LMAN-RA site pairs in 3 birds (represented by colored lines). **, p<0.005, LMAN-RA cross-covariance (trained minus baseline) modeled with fixed effects for intercept and syllable type and random effect for intercept and syllable type grouped by experiment ID. (B) Summary of change in LMAN and RA firing rates during training for targeted and non-targeted syllables. Firing rates, computed as mean spikes per second in the same premotor windows used for computing LMAN-RA cross-covariance, in the training period were normalized by dividing the firing rates at baseline. Each datapoint corresponds to spikes for a single syllable and recording site (RA: N = 29 Target, 152 Nontarget; LMAN: 14 Target, 64 Nontarget; across 11 experiments in 3 birds). n.s., not significant, t-test.

**Supplemental Figure 3.**
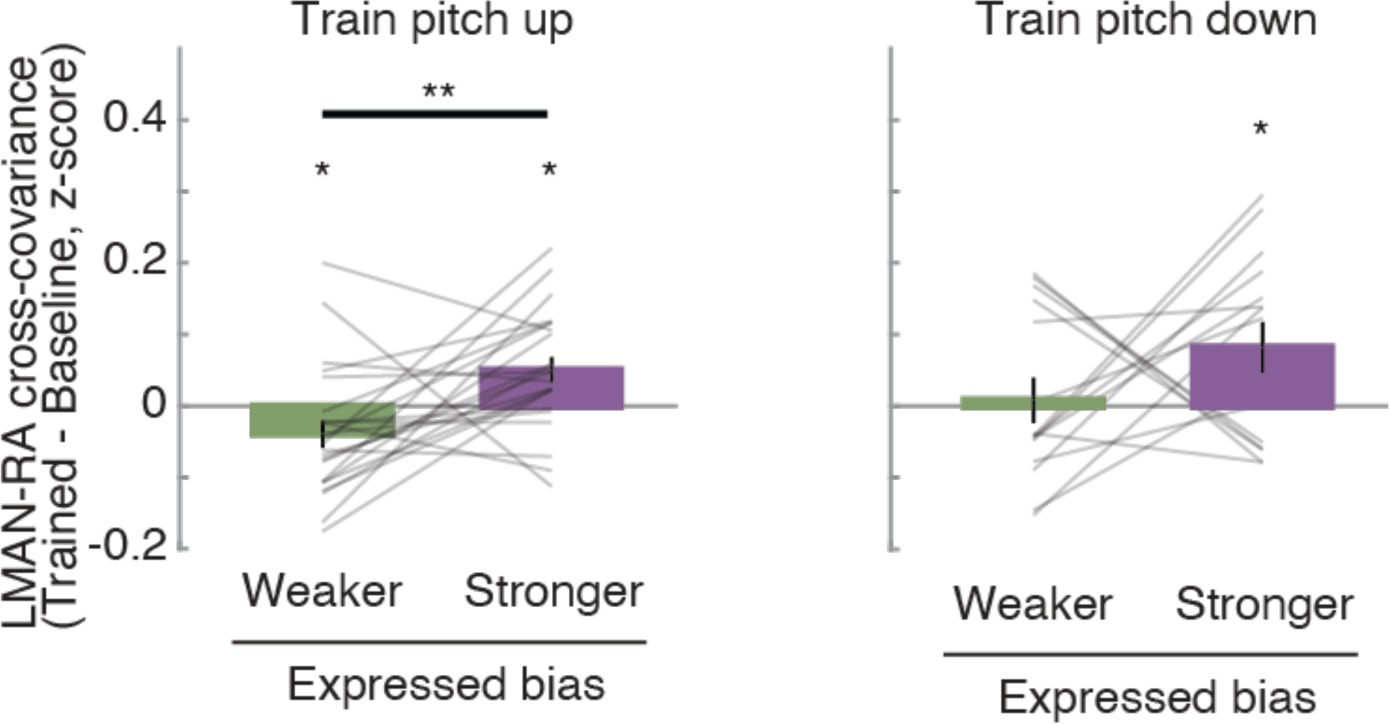
The strength of LMAN-RA co-fluctuations and the strength of adaptive motor bias are associated on a rendition-by-rendition basis, for both experiments training upwards or downwards pitch shifts (Related to Figure 3). For experiments training pitch up (left) or down (right), each panel shows change in LMAN-RA cross-covariance during training (Trained - Baseline), measured separately for renditions expressing “stronger” or “weaker” learning (up training: N = 23 LMAN-RA site pairs in 6 experiments in 3 birds; down training: 15 LMAN-RA site pairs in 5 experiments in 2 birds). *, **, p<0.05, p<0.005, Wilcoxon signed-rank test.

**Supplemental Figure 4.**
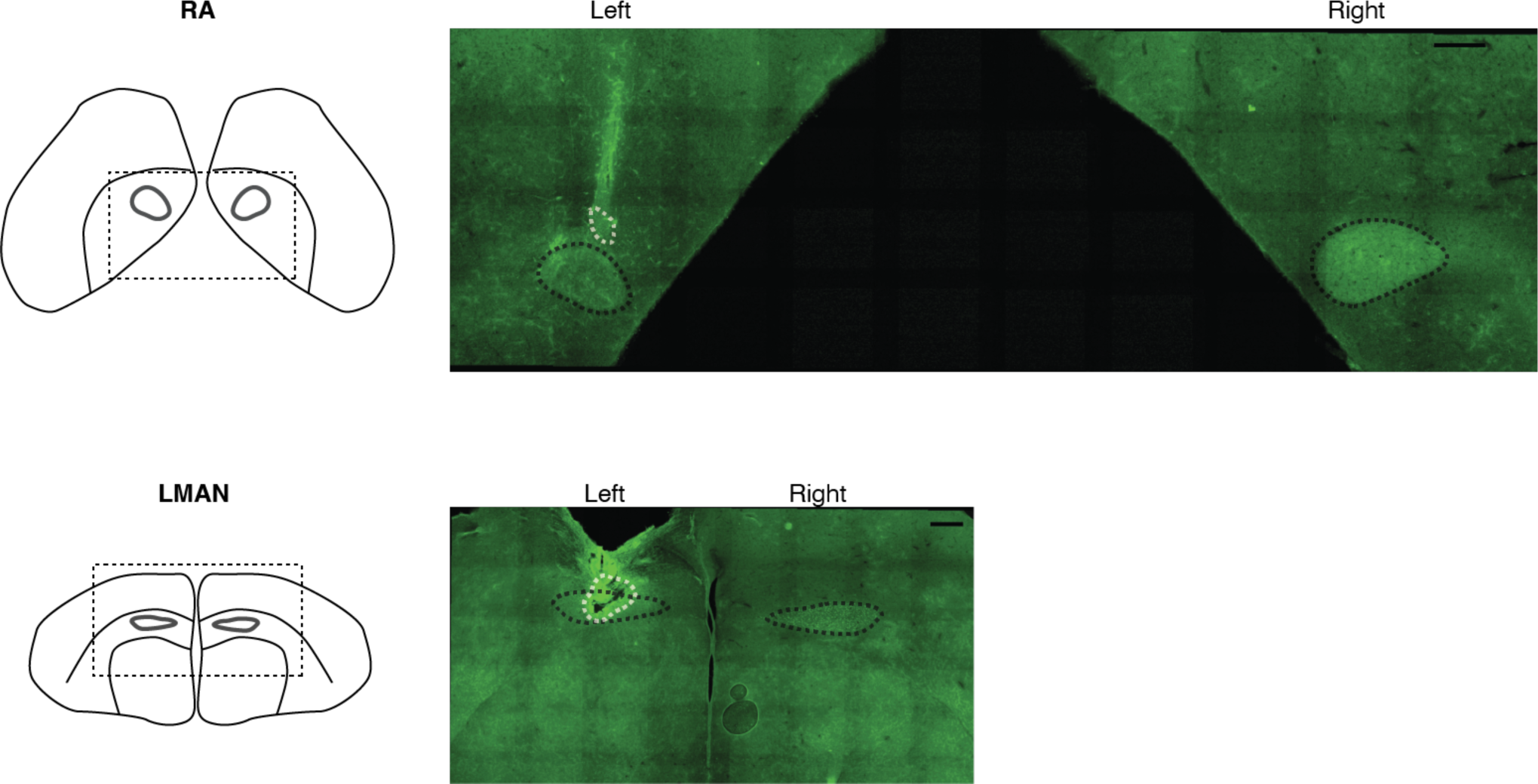
Histological verification of electrode locations. Post-mortem histology was used to confirm locations of electrodes for recording and stimulation experiments (see Methods). Example microimage of brain section stained against calcitonin gene related peptide (green), showing locations of recording electrode tips (white circles, marked by lesion) dorsal to LMAN and RA (black circles). The location of electrodes in this image confirms that electrodes entered LMAN and RA when lowered during experiments.

## References

1. Ali, F., Otchy, T.M., Pehlevan, C., Fantana, A.L., Burak, Y., Ölveczky, B.P., 2013. The Basal Ganglia Is Necessary for Learning Spectral, but Not Temporal, Features of Birdsong. Neuron 80, 494–506. https://doi.org/10.1016/j.neuron.2013.07.049

2. Amberg, G., Lindefors, N., 1989. Intracerebral microdialysis: II. mathematical studies of diffusion kinetics. Journal of Pharmacological Methods 22, 157–183. https://doi.org/10.1016/0160-5402(89)90012-0

3. Andalman, A.S., Fee, M.S., 2009. A basal ganglia-forebrain circuit in the songbird biases motor output to avoid vocal errors. Proceedings of the National Academy of Sciences 106, 12518–12523. https://doi.org/10.1073/pnas.0903214106

4. Aronov, D., Andalman, A.S., Fee, M.S., 2008. A Specialized Forebrain Circuit for Vocal Babbling in the Juvenile Songbird. Science 320, 630–634. https://doi.org/10.1126/science.1155140

5. Ashmore, R.C., Wild, J.M., Schmidt, M.F., 2005. Brainstem and Forebrain Contributions to the Generation of Learned Motor Behaviors for Song. J. Neurosci. 25, 8543–8554. https://doi.org/10.1523/JNEUROSCI.1668-05.2005

6. Bottjer, S., Miesner, E., Arnold, A., 1984. Forebrain lesions disrupt development but not maintenance of song in passerine birds. Science 224, 901–903. https://doi.org/10.1126/science.6719123

7. Bottjer, S.W., Halsema, K.A., Brown, S.A., Miesner, E.A., 1989. Axonal connections of a forebrain nucleus involved with vocal learning in zebra finches. J. Comp. Neurol. 279, 312–326. https://doi.org/10.1002/cne.902790211

8. Brainard, M.S., Doupe, A.J., 2000. Interruption of a basal ganglia–forebrain circuit prevents plasticity of learned vocalizations. Nature 404, 762–766. https://doi.org/10.1038/35008083

9. Brown, S.T., Raman, I.M., 2018. Sensorimotor Integration and Amplification of Reflexive Whisking by Well-Timed Spiking in the Cerebellar Corticonuclear Circuit. Neuron 99, 564–575.e2. https://doi.org/10.1016/j.neuron.2018.06.028

10. Charlesworth, J.D., Tumer, E.C., Warren, T.L., Brainard, M.S., 2011. Learning the microstructure of successful behavior. Nat Neurosci 14, 373–380. https://doi.org/10.1038/nn.2748

11. Charlesworth, J.D., Warren, T.L., Brainard, M.S., 2012. Covert skill learning in a cortical-basal ganglia circuit. Nature 486, 251–255. https://doi.org/10.1038/nature11078

12. Chi, Z., Margoliash, D., 2001. Temporal Precision and Temporal Drift in Brain and Behavior of Zebra Finch Song. Neuron 32, 899–910. https://doi.org/10.1016/S0896-6273(01)00524-4

13. Cowley, B.R., Snyder, A.C., Acar, K., Williamson, R.C., Yu, B.M., Smith, M.A., 2020. Slow Drift of Neural Activity as a Signature of Impulsivity in Macaque Visual and Prefrontal Cortex. Neuron 108, 551–567.e8. https://doi.org/10.1016/j.neuron.2020.07.021

14. Darshan, R., Wood, W.E., Peters, S., Leblois, A., Hansel, D., 2017. A canonical neural mechanism for behavioral variability. Nat Commun 8, 15415. https://doi.org/10.1038/ncomms15415

15. Dayan, E., Cohen, L.G., 2011. Neuroplasticity Subserving Motor Skill Learning. Neuron 72, 443–454. https://doi.org/10.1016/j.neuron.2011.10.008

16. Doya, K., Sejnowski, T.J., 1998. A Computational Model of Birdsong Learning by Auditory Experience and Auditory Feedback, in: Poon, P.W.F., Brugge, J.F. (Eds.), Central Auditory Processing and Neural Modeling. Springer US, Boston, MA, pp. 77–88. https://doi.org/10.1007/978-1-4615-5351-9_8

17. Doyon, J., Ungerleider, L., 2002. Functional anatomy of motor skill learning, in: Neuropsychology of Memory.

18. Fee, M.S., Goldberg, J.H., 2011. A hypothesis for basal ganglia-dependent reinforcement learning in the songbird. Neuroscience 198, 152–170. https://doi.org/10.1016/j.neuroscience.2011.09.069

19. Fee, M.S., Kozhevnikov, A.A., Hahnloser, R.H. r., 2004. Neural Mechanisms of Vocal Sequence Generation in the Songbird. Annals of the New York Academy of Sciences 1016, 153–170. https://doi.org/10.1196/annals.1298.022

20. Gadagkar, V., Puzerey, P.A., Chen, R., Baird-Daniel, E., Farhang, A.R., Goldberg, J.H., 2016. Dopamine neurons encode performance error in singing birds. Science 354, 1278–1282. https://doi.org/10.1126/science.aah6837

21. Giret, N., Kornfeld, J., Ganguli, S., Hahnloser, R.H.R., 2014. Evidence for a causal inverse model in an avian cortico-basal ganglia circuit. PNAS 111, 6063–6068. https://doi.org/10.1073/pnas.1317087111

22. Hahnloser, R.H.R., Kozhevnikov, A.A., Fee, M.S., 2006. Sleep-Related Neural Activity in a Premotor and a Basal-Ganglia Pathway of the Songbird. J Neurophysiol 96, 794–812. https://doi.org/10.1152/jn.01064.2005

23. Hessler, N.A., Doupe, A.J., 1999. Singing-Related Neural Activity in a Dorsal Forebrain–Basal Ganglia Circuit of Adult Zebra Finches. J. Neurosci. 19, 10461–10481.

24. Hikosaka, O., Nakamura, K., Nakahara, N., 2006. Basal Ganglia Orient Eyes to Reward. Journal of Neurophysiology 95, 567–584. https://doi.org/10.1152/jn.00458.2005

25. Hikosaka, O., Nakamura, K., Sakai, K., Nakahara, H., 2002. Central mechanisms of motor skill learning. Current Opinion in Neurobiology 12, 217–222. https://doi.org/10.1016/S0959-4388(02)00307-0

26. Hoffmann, L.A., Sober, S.J., 2014. Vocal Generalization Depends on Gesture Identity and Sequence. Journal of Neuroscience 34, 5564–5574. https://doi.org/10.1523/JNEUROSCI.5169-13.2014

27. Howard, I.S., Ingram, J.N., Franklin, D.W., Wolpert, D.M., 2012. Gone in 0.6 Seconds: The Encoding of Motor Memories Depends on Recent Sensorimotor States. Journal of Neuroscience 32, 12756– 12768. https://doi.org/10.1523/JNEUROSCI.5909-11.2012

28. Ignashchenkova, A., Dicke, P.W., Haarmeier, T., Thier, P., 2004. Neuron-specific contribution of the superior colliculus to overt and covert shifts of attention. Nat Neurosci 7, 56–64. https://doi.org/10.1038/nn1169

29. Jaffe, P.I., Brainard, M.S., 2020. Acetylcholine acts on songbird premotor circuitry to invigorate vocal output. eLife 9, e53288. https://doi.org/10.7554/eLife.53288

30. Kao, M.H., Brainard, M.S., 2006. Lesions of an Avian Basal Ganglia Circuit Prevent Context- Dependent Changes to Song Variability. Journal of Neurophysiology 96, 1441–1455. https://doi.org/10.1152/jn.01138.2005

31. Kao, M.H., Doupe, A.J., Brainard, M.S., 2005. Contributions of an avian basal ganglia–forebrain circuit to real-time modulation of song. Nature 433, 638–643. https://doi.org/10.1038/nature03127

32. Kao, M.H., Wright, B.D., Doupe, A.J., 2008. Neurons in a Forebrain Nucleus Required for Vocal Plasticity Rapidly Switch between Precise Firing and Variable Bursting Depending on Social Context. J. Neurosci. 28, 13232–13247. https://doi.org/10.1523/JNEUROSCI.2250-08.2008

33. Kawai, R., Markman, T., Poddar, R., Ko, R., Fantana, A.L., Dhawale, A.K., Kampff, A.R., Ölveczky, B.P., 2015. Motor cortex is required for learning but not for executing a motor skill. Neuron 86, 800– 812.

34. Kawashima, T., Zwart, M.F., Yang, C.-T., Mensh, B.D., Ahrens, M.B., 2016. The Serotonergic System Tracks the Outcomes of Actions to Mediate Short-Term Motor Learning. Cell 167, 933–946.e20. https://doi.org/10.1016/j.cell.2016.09.055

35. Kearney, M.G., Warren, T.L., Hisey, E., Qi, J., Mooney, R., 2019. Discrete Evaluative and Premotor Circuits Enable Vocal Learning in Songbirds. Neuron 104, 559–575.e6. https://doi.org/10.1016/j.neuron.2019.07.025

36. Kimpo, R.R., Theunissen, F.E., Doupe, A.J., 2003. Propagation of Correlated Activity through Multiple Stages of a Neural Circuit. J. Neurosci. 23, 5750–5761. https://doi.org/10.1523/JNEUROSCI.23-13-05750.2003

37. Kleim, J.A., 2004. Cortical Synaptogenesis and Motor Map Reorganization Occur during Late, But Not Early, Phase of Motor Skill Learning. Journal of Neuroscience 24, 628–633. https://doi.org/10.1523/JNEUROSCI.3440-03.2004

38. Knudsen, E., 1994. Supervised learning in the brain. J. Neurosci. 14, 3985–3997. https://doi.org/10.1523/JNEUROSCI.14-07-03985.1994

39. Kojima, S., Kao, M.H., Doupe, A.J., 2013. Task-related “cortical” bursting depends critically on basal ganglia input and is linked to vocal plasticity. PNAS 110, 4756–4761. https://doi.org/10.1073/pnas.1216308110

40. Kojima, S., Kao, M.H., Doupe, A.J., Brainard, M.S., 2018. The Avian Basal Ganglia Are a Source of Rapid Behavioral Variation That Enables Vocal Motor Exploration. J. Neurosci. 38, 9635–9647. https://doi.org/10.1523/JNEUROSCI.2915-17.2018

41. Koralek, A.C., Costa, R.M., Carmena, J.M., 2013. Temporally Precise Cell-Specific Coherence Develops in Corticostriatal Networks during Learning. Neuron 79, 865–872. https://doi.org/10.1016/j.neuron.2013.06.047

42. Koralek, A.C., Jin, X., Long II, J.D., Costa, R.M., Carmena, J.M., 2012. Corticostriatal plasticity is necessary for learning intentional neuroprosthetic skills. Nature 483, 331–335. https://doi.org/10.1038/nature10845

43. Kubota, M., Saito, N., 1991. NMDA receptors participate differentially in two different synaptic inputs in neurons of the zebra finch robust nucleus of the archistriatum in vitro. Neuroscience Letters 125, 107–109. https://doi.org/10.1016/0304-3940(91)90002-B

44. Kumar, A., Rotter, S., Aertsen, A., 2010. Spiking activity propagation in neuronal networks: reconciling different perspectives on neural coding. Nat Rev Neurosci 11, 615–627. https://doi.org/10.1038/nrn2886

45. Lemke, S.M., Ramanathan, D.S., Guo, L., Won, S.J., Ganguly, K., 2019. Emergent modular neural control drives coordinated motor actions. Nat Neurosci 22, 1122–1131. https://doi.org/10.1038/s41593-019-0407-2

46. Leonardo, A., Fee, M.S., 2005. Ensemble Coding of Vocal Control in Birdsong. J. Neurosci. 25, 652– 661. https://doi.org/10.1523/JNEUROSCI.3036-04.2005

47. Lipkind, D., Zai, A.T., Hanuschkin, A., Marcus, G.F., Tchernichovski, O., Hahnloser, R.H.R., 2017. Songbirds work around computational complexity by learning song vocabulary independently of sequence. Nat Commun 8, 1247. https://doi.org/10.1038/s41467-017-01436-0

48. Makino, H., Ren, C., Liu, H., Kim, A.N., Kondapaneni, N., Liu, X., Kuzum, D., Komiyama, T., 2017. Transformation of Cortex-wide Emergent Properties during Motor Learning. Neuron 94, 880–890.e8. https://doi.org/10.1016/j.neuron.2017.04.015

49. Margoliash, D., Yu, A.C., 1996. Temporal Heirarchical Control of Singing in Birds.

50. McCasland, J.S., 1987. Neuronal control of bird song production. The Journal of neuroscience 7, 23– 39.

51. Medina, J.F., Carey, M.R., Lisberger, S.G., 2005. The Representation of Time for Motor Learning. Neuron 45, 157–167. https://doi.org/10.1016/j.neuron.2004.12.017

52. Medina, J.F., Nores, W.L., Ohyama, T., Mauk, M.D., 2000. Mechanisms of cerebellar learning suggested by eyelid conditioning. Current Opinion in Neurobiology 8.

53. Mooney, R., Konishi, M., 1991. Two distinct inputs to an avian song nucleus activate different glutamate receptor subtypes on individual neurons. PNAS 88, 4075–4079. https://doi.org/10.1073/pnas.88.10.4075

54. Moorman, S., Ahn, J.-R., Kao, M.H., 2021. Plasticity of stereotyped birdsong driven by chronic manipulation of cortical-basal ganglia activity. Current Biology 31, 2619–2632.e4. https://doi.org/10.1016/j.cub.2021.04.030

55. Muller, J.R., Philiastides, M.G., Newsome, W.T., 2005. Microstimulation of the superior colliculus focuses attention without moving the eyes. Proceedings of the National Academy of Sciences 102, 524–529. https://doi.org/10.1073/pnas.0408311101

56. Narayanan, N.S., Laubach, M., 2006. Top-Down Control of Motor Cortex Ensembles by Dorsomedial Prefrontal Cortex. Neuron 52, 921–931. https://doi.org/10.1016/j.neuron.2006.10.021

57. Nordeen, K.W., Nordeen, E.J., 2010. Deafening-Induced Vocal Deterioration in Adult Songbirds Is Reversed by Disrupting a Basal Ganglia-Forebrain Circuit. Journal of Neuroscience 30, 7392–7400. https://doi.org/10.1523/JNEUROSCI.6181-09.2010

58. Nottebohm, F., Paton, J.A., Kelley, D.B., 1982. Connections of vocal control nuclei in the canary telencephalon. J. Comp. Neurol. 207, 344–357. https://doi.org/10.1002/cne.902070406

59. Noudoost, B., Chang, M.H., Steinmetz, N.A., Moore, T., 2010. Top-down control of visual attention. Current Opinion in Neurobiology 20, 183–190. https://doi.org/10.1016/j.conb.2010.02.003

60. Olveczky, B.P., Andalman, A.S., Fee, M.S., 2005. Vocal experimentation in the juvenile songbird requires a basal ganglia circuit. PLoS Biol. 3, e153. https://doi.org/10.1371/journal.pbio.0030153

61. Palmer, S.E., Wright, B.D., Doupe, A.J., Kao, M.H., 2021. Variable but not random: temporal pattern coding in a songbird brain area necessary for song modification. Journal of Neurophysiology 125, 540–555. https://doi.org/10.1152/jn.00034.2019

62. Perich, M.G., Gallego, J.A., Miller, L.E., 2018. A Neural Population Mechanism for Rapid Learning. Neuron 100, 964–976.e7. https://doi.org/10.1016/j.neuron.2018.09.030

63. Perkel, D.H., Gerstein, G.L., Moore, G.P., 1967. Neuronal Spike Trains and Stochastic Point Processes. Biophysical Journal 7, 419–440.

64. Ravbar, P., Lipkind, D., Parra, L.C., Tchernichovski, O., 2012. Vocal Exploration Is Locally Regulated during Song Learning. J. Neurosci. 32, 3422–3432. https://doi.org/10.1523/JNEUROSCI.3740-11.2012

65. Riehle, A., 1997. Spike Synchronization and Rate Modulation Differentially Involved in Motor Cortical Function. Science 278, 1950–1953. https://doi.org/10.1126/science.278.5345.1950

66. Rochet-Capellan, A., Ostry, D.J., 2011. Simultaneous Acquisition of Multiple Auditory-Motor Transformations in Speech. Journal of Neuroscience 31, 2657–2662. https://doi.org/10.1523/JNEUROSCI.6020-10.2011

67. Rochet-Capellan, A., Richer, L., Ostry, D.J., 2012. Nonhomogeneous transfer reveals specificity in speech motor learning. Journal of Neurophysiology 107, 1711–1717. https://doi.org/10.1152/jn.00773.2011

68. Rueda-Orozco, P.E., Robbe, D., 2015. The striatum multiplexes contextual and kinematic information to constrain motor habits execution. Nature Neuroscience 18, 453–460. https://doi.org/10.1038/nn.3924

69. Sawada, M., Kato, K., Kunieda, T., Mikuni, N., Miyamoto, S., Onoe, H., Isa, T., Nishimura, Y., 2015. Function of the nucleus accumbens in motor control during recovery after spinal cord injury. Science 350, 98–101. https://doi.org/10.1126/science.aab3825

70. Scharff, C., Nottebohm, F., 1991. A comparative study of the behavioral deficits following lesions of various parts of the zebra finch song system: implications for vocal learning. J. Neurosci. 11, 2896– 2913.

71. Semedo, J.D., Zandvakili, A., Machens, C.K., Yu, B.M., Kohn, A., 2019. Cortical Areas Interact through a Communication Subspace. Neuron 102, 249–259.e4. https://doi.org/10.1016/j.neuron.2019.01.026

72. Shadmehr, R., Holcomb, H.H., 1997. Neural correlates of motor memory consolidation. Science 277, 821–825.

73. Sober, S.J., Wohlgemuth, M.J., Brainard, M.S., 2008. Central Contributions to Acoustic Variation in Birdsong. J. Neurosci. 28, 10370–10379. https://doi.org/10.1523/JNEUROSCI.2448-08.2008

74. Stavisky, S.D., Kao, J.C., Ryu, S.I., Shenoy, K.V., 2017. Trial-by-Trial Motor Cortical Correlates of a Rapidly Adapting Visuomotor Internal Model. J. Neurosci. 37, 1721–1732. https://doi.org/10.1523/JNEUROSCI.1091-16.2016

75. Stepanek, L., Doupe, A.J., 2010. Activity in a cortical-basal ganglia circuit for song is required for social context-dependent vocal variability. J. Neurophysiol. 104, 2474–2486. https://doi.org/10.1152/jn.00977.2009

76. Tchernichovski, O., Mitra, P.P., Lints, T., Nottebohm, F., 2001. Dynamics of the Vocal Imitation Process: How a Zebra Finch Learns Its Song. Science 291, 2564–2569. https://doi.org/10.1126/science.1058522

77. Tian, L.Y., Brainard, M.S., 2017. Discrete Circuits Support Generalized versus Context-Specific Vocal Learning in the Songbird. Neuron 96, 1168–1177.e5. https://doi.org/10.1016/j.neuron.2017.10.019

78. Troyer, T.W., Bottjer, S.W., 2001. Birdsong: models and mechanisms. Current Opinion in Neurobiology 11, 721–726. https://doi.org/10.1016/S0959-4388(01)00275-6

79. Troyer, T.W., Doupe, A.J., 2000a. An Associational Model of Birdsong Sensorimotor Learning II. Temporal Hierarchies and the Learning of Song Sequence. J Neurophysiol 84, 1224–1239.

80. Troyer, T.W., Doupe, A.J., 2000b. An Associational Model of Birdsong Sensorimotor Learning I. Efference Copy and the Learning of Song Syllables. Journal of Neurophysiology 84, 1204–1223. https://doi.org/10.1152/jn.2000.84.3.1204

81. Tumer, E.C., Brainard, M.S., 2007. Performance variability enables adaptive plasticity of ‘crystallized’ adult birdsong. Nature 450, 1240–1244. https://doi.org/10.1038/nature06390

82. Turner, R.S., Desmurget, M., 2010. Basal ganglia contributions to motor control: a vigorous tutor. Current Opinion in Neurobiology 20, 704–716. https://doi.org/10.1016/j.conb.2010.08.022

83. Veuthey, T.L., Derosier, K., Kondapavulur, S., Ganguly, K., 2020. Single-trial cross-area neural population dynamics during long-term skill learning. Nat Commun 11, 4057. https://doi.org/10.1038/s41467-020-17902-1

84. Vu, E.T., Mazurek, M.E., Kuo, Y.C., 1994. Identification of a forebrain motor programming network for the learned song of zebra finches. J. Neurosci. 14, 6924–6934.

85. Wagner, M.J., Kim, T.H., Kadmon, J., Nguyen, N.D., Ganguli, S., Schnitzer, M.J., Luo, L., 2019. Shared Cortex-Cerebellum Dynamics in the Execution and Learning of a Motor Task. Cell 177, 669–682.e24. https://doi.org/10.1016/j.cell.2019.02.019

86. Wainscott, S.K., 2004. Internal Models and Contextual Cues: Encoding Serial Order and Direction of Movement. Journal of Neurophysiology 93, 786–800. https://doi.org/10.1152/jn.00240.2004

87. Warren, T.L., Tumer, E.C., Charlesworth, J.D., Brainard, M.S., 2011. Mechanisms and time course of vocal learning and consolidation in the adult songbird. Journal of Neurophysiology 106, 1806–1821. https://doi.org/10.1152/jn.00311.2011

88. Woolley, S.C., Rajan, R., Joshua, M., Doupe, A.J., 2014. Emergence of Context-Dependent Variability across a Basal Ganglia Network. Neuron 82, 208–223. https://doi.org/10.1016/j.neuron.2014.01.039

89. Zandvakili, A., Kohn, A., 2015. Coordinated Neuronal Activity Enhances Corticocortical Communication. Neuron 87, 827–839. https://doi.org/10.1016/j.neuron.2015.07.026

